# Sex-linked gene traffic underlies the acquisition of sexually dimorphic UV color vision in *Heliconius* butterflies

**DOI:** 10.1101/2022.07.04.498748

**Authors:** Mahul Chakraborty, Angelica Guadalupe Lara, Andrew Dang, Kyle J. McCulloch, Dylan Rainbow, David Carter, Luna Thanh Ngo, Edwin Solares, Iskander Said, Russ Corbett-Detig, Lawrence E. Gilbert, J.J. Emerson, Adriana D. Briscoe

## Abstract

Butterflies have photoreceptor cells that are sensitive to the ultraviolet part of the spectrum due to *ultraviolet-sensitive rhodopsin* (*UVRh*), a gene that has been duplicated in the *Heliconius* genus. In individuals expressing UVRh1 and UVRh2, electrophysiological and behavioral studies demonstrate that these opsin proteins enable discrimination of UV wavelengths. This behavioral trait varies between species, being absent in *H. melpomene* and limited to females in *H. erato*. To identify the evolutionary origins of this trait, we first examined UV color vision in *H. charithonia*, a species related to *H. erato* in the *sara/sapho* group. We found that this species also has sexually dimorphic UV color vision. To identify the genetic basis of this trait, we built a reference-grade genome assembly of *H. charithonia*. We discovered that one duplicate, *UVRh1*, is present on the W chromosome, making it obligately female-specific. We employed gDNA PCR assays of *UVRh1* across the *Heliconius* genus. In species with sexually dimorphic *UVRh1* mRNA expression, *UVRh1* gDNA is absent in males, whereas in species with sexually monomorphic *UVRh1* mRNA expression, *UVRh1* gDNA is found in both sexes. The presence or absence of male *UVRh1* expression across the *Heliconius* phylogeny supports a model where sexual dimorphism was acquired early via movement of a gene duplication to the W-chromosome. We used CRISPR-Cas9 to engineer a deletion in the *UVRh1* locus in female *H. charithonia* and use immunohistochemistry to show that UVRh1 protein expression is absent in mutant tissue, similar to that of males. Our results show that a rare behavioral phenotype, sex-specific UV color vision, was acquired via sex chromosome gene traffic of a duplicated UV rhodopsin.

---

The acquisition of novel sexually dimorphic traits poses an evolutionary puzzle: how does a new trait arise and how does it become sex-limited? The persistence of sexual dimorphism over evolutionary timescales implies that optimal phenotypes for such traits differ between sexes. Discovering the molecular events leading to different phenotypic outcomes is crucial to understanding the evolutionary mechanisms that resolve such sexually-mediated tradeoffs (Rice 1984; Fry 2010). Candidate mechanisms include: the “pleiotropy-mechanism” (PM) whereby the sex-limitation and the new trait arise simultaneously, avoiding a tradeoff; and the “modifier-mechanism” (MM), whereby a new monomorphic trait arises first, followed by the acquisition of modifiers that restore the ancestral state in one sex, thereby resolving the tradeoff (Turner 1978; Rice 1984). The visual system offers one useful model for understanding this process. The genetics and physiology of vision is well-understood for a number of animals, and many instances of sexual dimorphism in the visual system, specifically in the expression of opsins or photostable filtering pigments in insects, have been documented (Arikawa et al. 2005; Sison-Mangus et al. 2006; Ogawa et al. 2012; Perry and Desplan 2016; McCulloch et al. 2017; Liénard et al. 2021). Furthermore, sexual dimorphism for color vision behavior is observed in both New World monkeys (Jacobs 1998) and in the butterfly genus *Heliconius* (Finkbeiner and Briscoe 2021).

In animal vision, distinct photoreceptor cell subtypes can be sensitive to different wavelengths of light. Variation in color sensitivity is primarily conferred by differences in the rhodopsin pigments–opsin proteins together with a chromophore–that absorb light. The integration of neural signals from different photoreceptor cells leads to color vision. In the genus *Heliconius*, there are four opsin genes, which encode a green wavelength-absorbing (LWRh), a blue wavelength-absorbing (BRh), and two ultraviolet wavelength-absorbing (UVRh1 and UVRh2) rhodopsins. The two UV rhodopsins are the consequence of a recent gene duplication that occurred ∼18.5 million years ago in the ancestor of all *Heliconius* butterflies (Briscoe et al. 2010; Kozak et al. 2015). Individuals expressing both UVRh1 and UVRh2 opsins can have at least two distinct ultraviolet-sensitive photoreceptor cell types, suggesting that these individuals can distinguish different UV wavelengths. Indeed, intracellular recordings have demonstrated different spectral sensitivities for two UV cell types in *H. erato* females (McCulloch et al. 2016). Behavioral analysis has further shown that female *H. erato* butterflies can distinguish different UV wavelengths (Finkbeiner and Briscoe 2021). On the other hand, *H. melpomene* lacks this type of UV photoreceptor dimorphism and UV color vision behavior (Finkbeiner and Briscoe 2021; McCulloch et al. 2022). Despite extensive genomic work in the *Heliconius* genus, including a reference genome for *H. melpomene* (Davey et al. 2016), the *erato/sara/sapho* clade lacks a genome assembly placing *UVRh1* on its chromosome (Lewis et al. 2016), which is crucial to understanding the evolution of sexually dimorphic UV color vision.

To uncover the path evolution followed in acquiring divergent UV color vision phenotypes between the *erato*/*sara/sapho* and *doris/melpomene* clades (Fig. 1A), we needed to document the location, structure, and genomic context of both *UVRh* duplicates in representatives of both clades. To accomplish this for the *erato*/*sara*/*sapho* clade, we built a reference-quality genome assembly for *H. charithonia–*a species exhibiting differences in the flower types visited by males and females (Mendoza-Cuenca and Macías-Ordóñez 2005)–to compare against the existing high-quality draft genome of *H. melpomene* (Davey et al. 2016). We used long read sequencing and RNA-seq data to create and annotate a highly contiguous, complete, and accurate reference-grade genome assembly (Figs. S1-4). In addition to recovering 99% of Lepidopteran Benchmarking Universal Single Copy Orthologs (BUSCO) in the assembly (Manni et al. 2021), 50% of the sequence is represented by contigs 16.4 Mb and longer (i.e. contig N50 = 16.4 Mb). Upon scaffolding with Hi-C, we attained sequences that span chromosomes nearly end-to-end (scaffold N50 = 17 Mb). Moreover, we also recovered a large scaffold representing the W-chromosome (Fig 1B-E). To our surprise, *UVRh1* is located on the W scaffold, in contrast with *H. melpomene*, in which both *UVRh* duplicates are autosomal (Fig. 2A-E)(*Heliconius* Genome Consortium 2012). In the outgroup species *Eueides isabella* and *Dryas iulia*, a single *UVRh* gene occupies the genomic location corresponding to *Heliconius UVRh2* (Fig. 2E).

**Figure 1.**
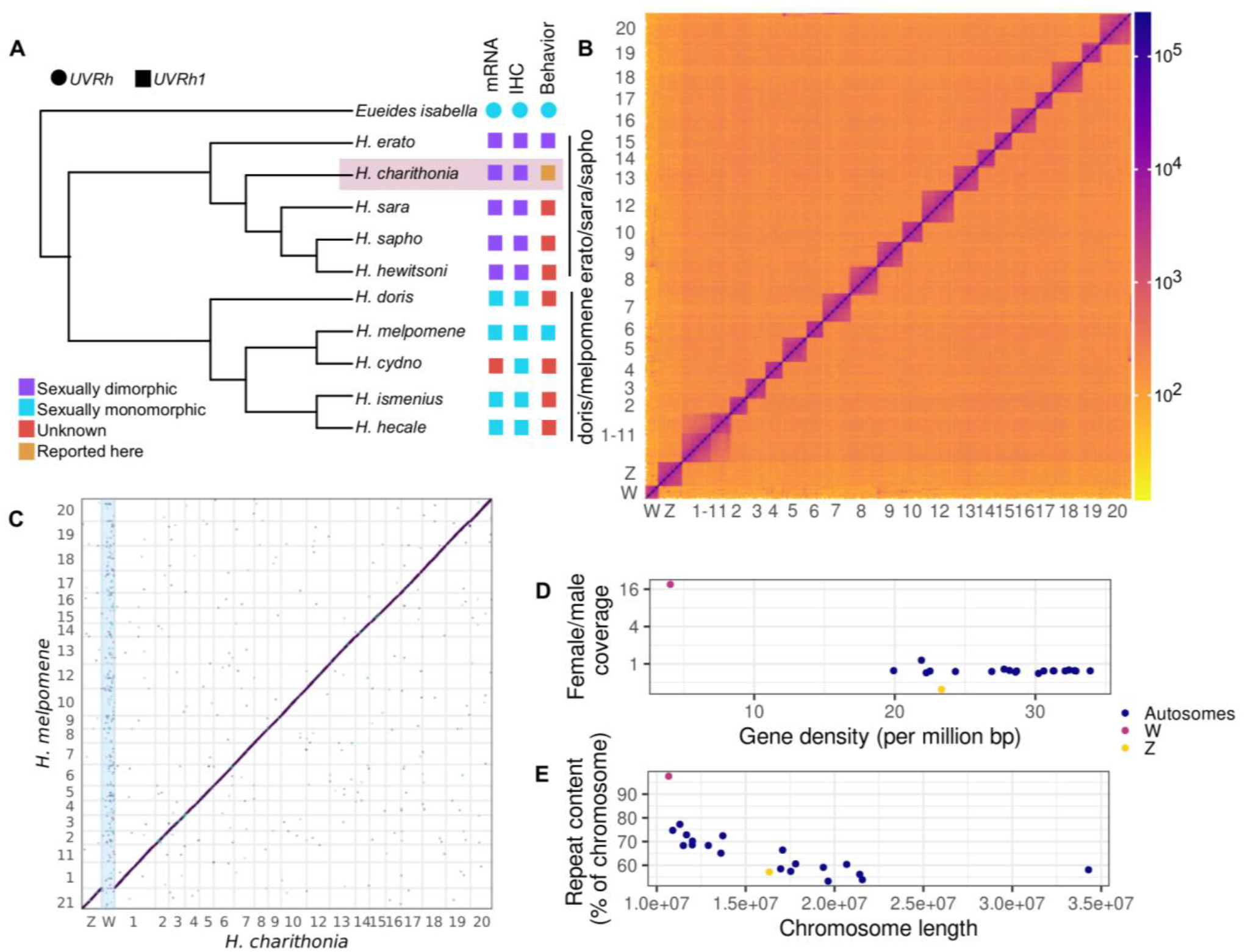
A *de novo* genome assembly of *Heliconius charithonia* and its phylogenetic relationship with species showing sexually monomorphic and dimorphic *UVRh1* expression. A) A cladogram showing the phylogenetic relationship among 10 *Heliconius* species, including *H. charithonia* and outgroup species *E. isabella*, based on Kozak et al. (2015). Five species from the *erato*/*sara/sapho* clades show sexually dimorphic expression of *UVRh1* mRNA and protein (immunohistochemistry or IHC) and female *H. erato* show UV color vision behavior. UV color discrimination in *H. charithonia* is reported in the present study. UVRh1 expression in other species is either sexually monomorphic or unknown. B) A Hi-C contact density map of the *H. charithonia* genome assembly showing 21 chromosomes. Chromosome 1 is a fusion of two chromosomes. C) An alignment dot plot between the genome assemblies of *H. melpomene* and *H. charithonia*. As shown here, *H. charithonia* Chromosome 1 is a fusion of *H. Melpomene* Chromosomes 1 & 11 and the W scaffold has no corresponding sequence in the *H. melpomene* assembly, which represents a male genome. D) Gene density and the ratio of female and male short read coverage of 21 *H. charithonia* chromosomes. The W scaffold has very few protein coding genes and virtually no unique sequence shared with a male genome. E) Relationship between chromosome length and repeat content of *H. charithonia* chromosomes. The chromosomes show a negative correlation between length and repeat content.

**Figure 2.**
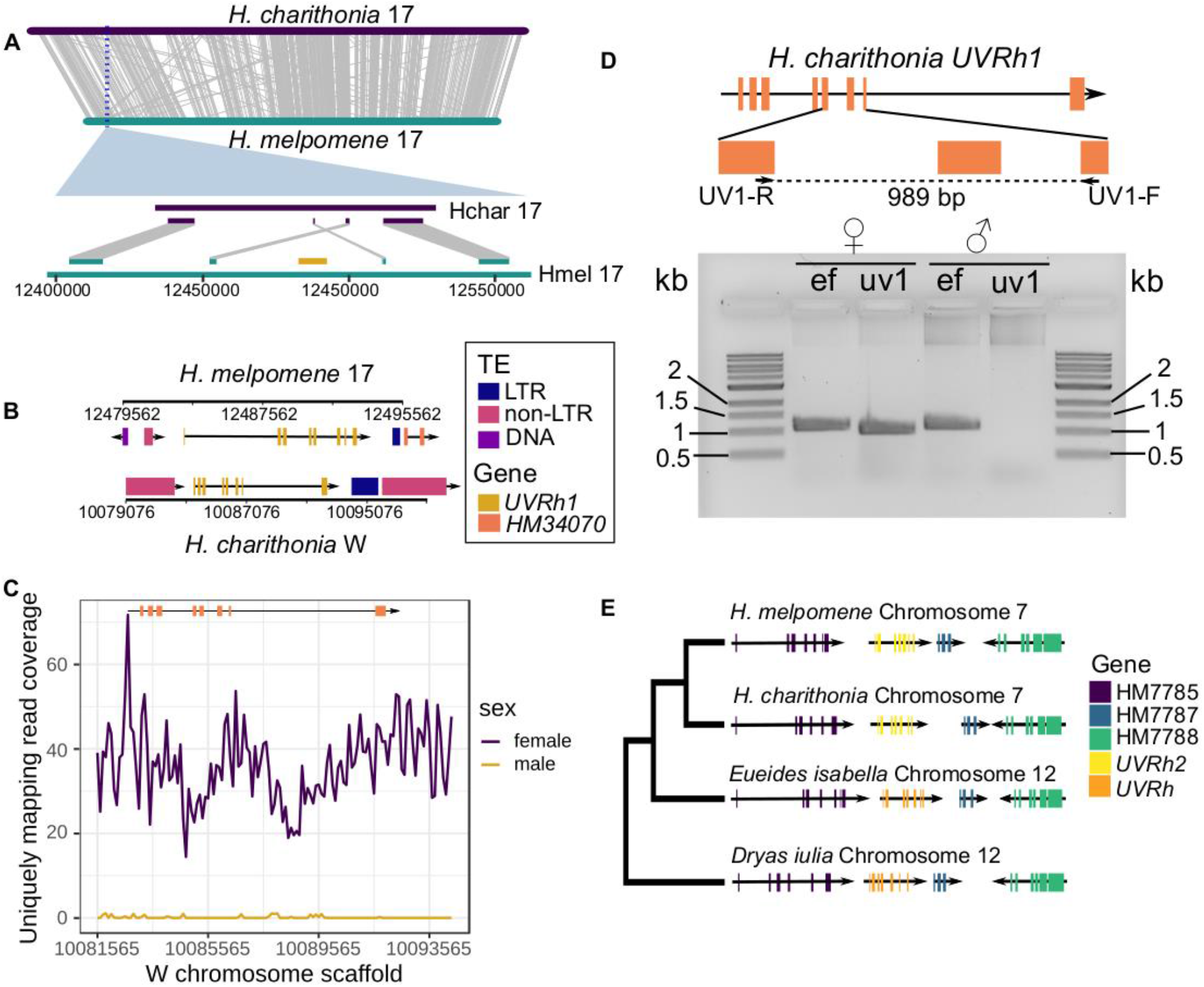
Genomic location of *UVRh1* and *UVRh2* in *Heliconius*. A) Alignment between *H. charithonia* and *H. melpomene* Chromosome 17 showing global synteny between the two chromosomes, although *UVRh1* is missing from *H. charithonia* Chromosome 17. B) *UVRh1* cDNA maps to the W scaffold in *H. charithonia* and shares the same number of exons and introns as *H. melponene UVRh1*. Presence of similar TE sequences on both sides of *UVRh1* in *H. melpomene* and *H. charithonia* indicates a possible role of TEs in translocation of *UVRh1*. C) Mapping coverage of uniquely mapping male and female Illumina paired end reads to the W scaffold region containing *UVRh1*. Virtually zero coverage of male reads supports the female linkage of *UVRh1*. D) Confirmation of W-linkage of *UVRh1* using PCR. A *UVRh1*-specific primer pair (uv1) amplifies only female *H. charithonia* gDNA but not male gDNA. The control primer *EF1a* (ef) amplifies both male and female gDNA. E) Genomic location of *UVRh2* in *H. Melpomene* and in *H. charithonia* and of *UVRh* in two outgroup species *Eueides isabella* and *Dryas iulia* (Cicconardi et al. 2021; Lewis et al. 2021) along with three other genes in *H. melpomene* reference genome release 2.5 (Davey et al. 2016). Conserved synteny of the genes suggest that *UVRh2*, on *Heliconius* chromosome 7, retains the genomic location of ancestral single copy *UVRh*, which is on *Eueides* chromosome 12.

The descendant of this ancestral locus resides on chromosome 12 in *E. isabella*, which is syntenic with the location of *UVRh2* on chromosome 7 in *H. melpomene*, while *UVRh1* is present on chromosome 17 in *H. melpomene* (Fig. 2A,E). To determine the sex linkage of *UVRh1* in representative species across the genus, we designed gDNA PCR assays targeting *UVRh1* in 10 species, five of which show sexually dimorphic UVRh1 protein expression (McCulloch et al. 2017). We successfully amplified and sequenced PCR products specific to *UVRh1* for all species (Fig. 3, S5, S6). For species in the *doris*/*melpomene* clades, we recovered *UVRh1* amplicons in both sexes. However, for species in the *erato*/*sara/sapho* clades, the *UVRh1* amplicons were limited to females. In all cases, positive control amplicons were present in both sexes (Fig. 3B). Using a phylogeny of 20 species and a maximum likelihood approach, we inferred that absence of *UVRh1* in males was the likely ancestral state of the *erato/sara/sapho* clade. However, we were unable to infer whether or not *UVRh1* was absent in males at the base of the genus *Heliconius* because the *erato/sara/sapho* and the *doris/melpomene* clades are sister groups (Fig. S8). Either *UVRh* was first duplicated onto the W-chromosome in the *Heliconius* common ancestor, limiting *UVRh1* to females, or *UVRh* was first duplicated onto an autosome. Under the first scenario, a translocation in the common ancestor of the *doris*/*melpomene* clades then moved *UVRh1* from the W to the homolog of chromosome 17 in *H. melpomene*, initiating autosomal linkage. Under the second scenario, a translocation in the common ancestor of the *erato/sara/sapho* clades then moved *UVRh1* from the homolog of chromosome 17 to the W.

**Figure 3.**
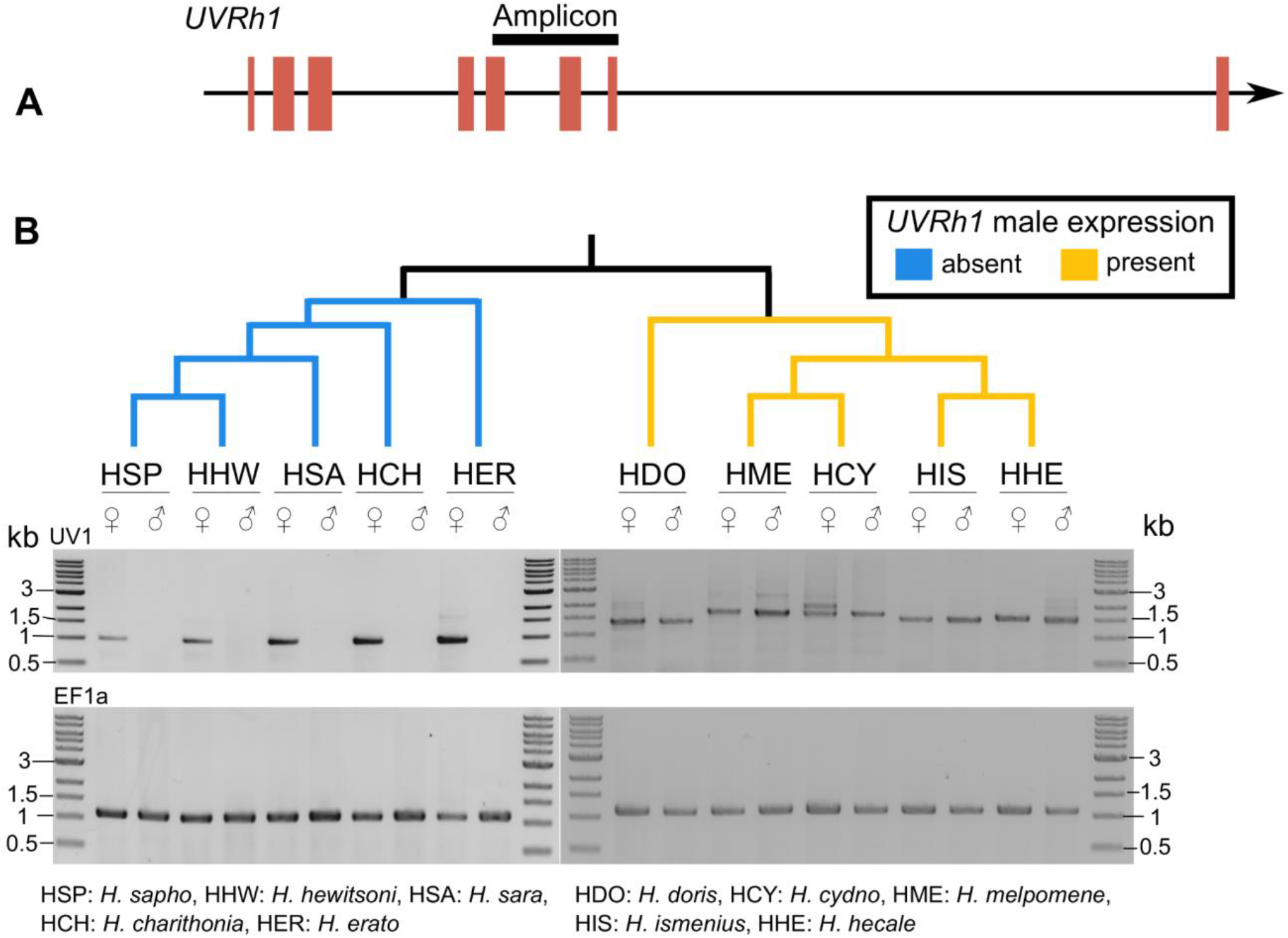
Determining *UVRh1* linkage across the genus *Heliconius* using gDNA PCR. A) Cartoon of the relative length of the *UVRh1* amplicon used to determine sex-linkage of *UVRh1* in 10 *Heliconius* species. B) *UVRh1* PCR products from 10 *Heliconius* species, five of which show sexually dimorphic *UVRh1* amplification. Only female DNA from the five species shown in blue and both sexes in the five species shown in yellow produced the *UVRh1* amplicon. *H. cydno* females produced an additional *UVRh1* PCR product that is absent in males (Fig. S5). The cladogram on top of the gel is based on the published *Heliconius* phylogeny (Kozak et al. 2015).

To establish that the *H. charithonia* gene we annotate as *UVRh1* encodes the UVRh1 protein in female photoreceptor cells, we knocked out the *UVRh1* gene in the adult eye. To achieve this via CRISPR-mediated deletion, we designed two guide RNAs targeting the 2nd and 3rd exons of *UVRh1*. We co-injected Cas9 and the gRNAs into 0-1 hour embryos and reared the survivors into adulthood. To visualize the locations of the short-wavelength opsins, the eyes were fixed and stained with anti-UVRh1, -UVRh2, and -BRh opsin antibodies. Adult CRISPR-edited female eye tissue exhibited mosaicism for two tissue types: female tissue with UVRh1, UVRh2, and BRh opsin-expressing photoreceptors, and male-like tissue containing only UVRh2 and BRh opsin-expressing photoreceptors (Fig. 4, Fig. S7J, S9).

**Figure 4.**
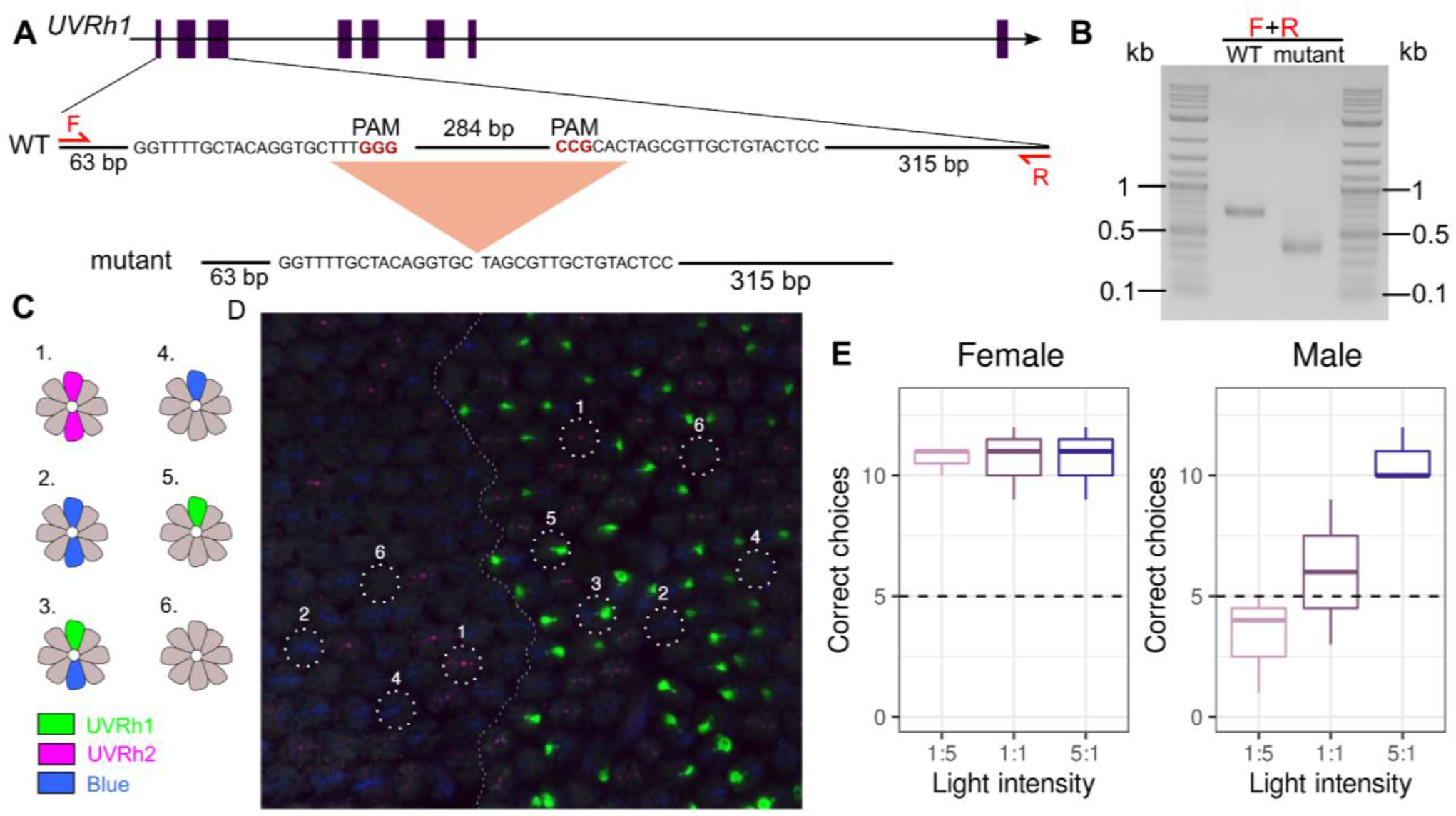
Targeted CRISPR/Cas9 knockout of *UVRh1* in an adult *H. charithonia* female eye and UV color vision behavioral trials. A) *UVRh1* gene model and sequence showing the location of a 296 bp deletion resulting from CRISPR/Cas 9 mutagenesis. B) PCR products of *UVRh1* genomic region flanking the deletion. C) Cartoon: Wild-type *H. charithonia* female retinas have at least six types of ommatidia based on opsin expression in the R1 and R2 photoreceptor cells: 1. UVRh2/UVRh2, 2. BRh/BRh, 3. UVRh1/BRh, 4. BRh-LWRh/BRh, 5. UVRh1/LWRh-BRh, 6. LWRh-BRh/LWRh-BRh (LWRh1 and BRh co-expression shown in Fig. S9). D) CRISPR targeted *UVRh1* produces adult female retinas that lack UVRh1 protein in large domains (middle panel (left), compared to wild-type, middle panel (right)). Knockout of *UVRh1* eliminates UVRh1 (green) protein expression in ommatidial types 3 and 5 (left) while UVRh2 (magenta) ommatidial type 1 and BRh (blue) ommatidial types 2 and 4 are retained (left). E) Number of correct choices by *H. charithonia* adult butterflies for the rewarded wavelength (+) when given a choice between 390 nm (+) and 380 nm (-) light under varying intensities. N=3 biological replicates per sex, N=15 choice trials per intensity. Females show a significant preference for the rewarded light over all light intensities (p-value=0.01) while males only show a significant preference for the rewarded light at the 5:1 intensity (p-value=0.01). Boxes represent upper and lower quartiles with median; whiskers indicate 25th and 75th percentiles.

Finally, we conducted behavioral trials to confirm that the expression of UVRh1 and UVRh2 in photoreceptor cells in female *H. charithonia* eyes confers the ability of adults to discriminate between different wavelengths of ultraviolet light. Adult male and female butterflies were trained to associate a sugar reward with 390 nm UV light following the protocol of Finkbeiner and Briscoe (2021). After training, adults were then given a choice between two UV lights: a rewarded light (390 nm), and an unrewarded light (380 nm)(Fig. S10). Individuals that flew to a light source were scored as selecting that light source. Females exhibited a strong and significant preference for 390 nm, the rewarded light, regardless of the relative intensity of the stimuli (z value=2.739, p-value=0.01) (Fig. 4E, Tables S2, S3), indicating that females have UV color vision. In contrast, males exhibited a preference for the brighter light source, correctly and significantly selecting the trained wavelength only when it was brightest (z value=2.739, p-value=0.01)(Fig. 4E, S8, Tables S2, S3), an indication of positive UV phototaxis but not UV color vision.

Here we show that gene traffic from an autosome to the W chromosome is the genetic mechanism behind the acquisition of sex-specific UV opsin expression in *Heliconius*. Relocation of the duplicate gene *UVRh1* to the W chromosome in the *erato*/*sara/sapho* clades leads to its absence in males, making it female-specific. A requirement for color vision is the specialization of photoreceptor cells to be sensitive to different wavelengths of light. Previous studies showed that rapid molecular divergence of *UVRh2* compared to *UVRh1* has led to extensive amino acid variation between the two duplicates (Briscoe et al. 2010), likely resulting in spectral tuning of UVRh2 associated with functional divergence in photoreceptor spectral sensitivity (McCulloch et al. 2016; McCulloch et al. 2022). Regardless of the specific path taken in UV opsin evolution, what is clear is that the duplication of the ancestral UV rhodopsin was followed by acquisition both of a novel expression pattern in females and of a novel protein function.

To classify the path followed in the molecular evolution of novel sexually dimorphic UV color vision, we consider the two previous models–the pleiotropy mechanism (PM) (Rice 1984) and the modifier model (MM) (Rice 1984)–and propose a third: partitioning-first (PF) whereby the genetic basis of the trait is first partitioned by sex, followed by a shift in the phenotype. In cases of duplication of genes like opsins, each copy can in principle correspond to independently mutable instances of the trait. This has two relevant consequences. First, gene duplication may avoid sexually antagonistic fitness tradeoffs, as independent fitness optima can be achieved simultaneously for each copy (Connallon and Clark 2011; Chakraborty and Fry 2015). Second, duplication permits sex-biased partitioning to precede the shift of a trait fitness value. For example, retrogenes successfully escaping the X chromosome in mammals immediately cease existing in an environment subject to meiotic X-inactivation, a process specific to male biology (Long and Emerson 2017). In the evolution of UV color vision, the path to the phenotype shift and the sex-specificity did not happen simultaneously so the pleiotropy model is a poor fit. Since most of the rapid amino acid evolution of UVRh2 occurred in a common ancestor of *Heliconius* with two *UVRh* genes, the order of the mutations will determine whether the spectral sensitivity shift (MM) or sexually dimorphic partitioning (PF) happened first. Finer genome-level sampling of *Heliconius* will facilitate more refined phylogenetic hypotheses (Turner 1976), potentially resolving the specific evolutionary sequence of events. It is intriguing too, that the *erato/sara/sapho* clade is united not only by the loss of *UVRh1* in males but also in pupal mating and its associated morphology (e.g. the absence of signa in female bursa copulatrix)(Penz 1999) and behavior (e.g. the ability of males to discriminate the sex of pupae)(Estrada et al. 2010). These traits may be intriguing candidates for driving differences in vision between the two major *Heliconius* subclades characterized here.

X-linked opsin gene expression has been shown to underlie sexual dimorphism of red-green color vision in New World monkeys (Hunt et al. 1998). However an important difference exists between the red-green color vision dimorphism of NW primates, which is based on a single-gene allelic system, and the UV color vision dimorphism in *Heliconius* described here. Elucidating the transcriptional mechanisms that control UV opsin expression will shed light into the processes regulating sex-specific gene expression, and the identification of associated downstream neural circuitry changes will provide insights into the evolution of behavioral differences between the sexes. In conclusion we show that an extreme form of female-limited UV color vision behavior in butterflies has evolved via gene duplication followed by sex chromosome translocation and that this finding reveals how novel sex-specific complex traits can arise in a short evolutionary time.

## Acknowledgements

We thank Javier Rodarte, Zachary Johnson, Aline Rangel Olguin, Yuan Tao, Furong Yuan, JP Lawrence, Matthew Aardema, Peter Andolfatto and Stephannie Seng for technical assistance and UCI’s Optical Biology Core for microscopy access. This work was funded in part by NSF grant IOS-1656260 to A.D.B, NIH grants R01GM123303-1 to J.J.E. and K99GM129411 to M.C., and through grants CA-62203 and GM-076516 supporting UCI’s Optical Biology Core Facility.

## Methods

### Butterfly samples

A single pair mating of *H. charithonia* was generated in the greenhouse at the University of Texas, Austin in October 2017. From this mating, a single adult female F1 specimen was used in the generation of Hi-C data. Extraction of high molecular weight from other F1 adults from this mating did not yield DNA of sufficiently high quality so in March 2018, a female pupa descended from the UT colony was used to generate the PacBio data. Two other male and female individuals from the same source were used for Illumina DNA short-read sequencing. Embryos used for CRISPR injection were collected from mated females descended from pupae sourced from the Costa Rica Entomological Supply. Locality information for specimens used in PCR and behavioral experiments are given in Table S4.

### DNA extraction and Sequencing

High molecular weight genomic DNA was extracted from a single *H. charithonia* female pupa following established protocols (Chakraborty et al. 2016). Briefly, the pupa was cut open from the posterior end using a razor blade and the soft tissue was squeezed out using a homogenizer. The tissue was homogenized in buffer G2 of Qiagen Blood and Cell Culture Tissue Kit and rest of the DNA extraction was carried out as described in Chakraborty et al. (2016). DNA was sheared with 10 plunges of 21 gauge blunt-end needle followed by 10 plunges of 24 gauge blunt-end needle. The sheared DNA was size selected on Blue Pippin using 20 kb minimum cut-off length and a library was created from this size selected DNA. The library was sequenced with 33 SMRTcells on the Pacific Biosciences RS II platform, producing a total of 49.5 Gbp sequences (50% of the reads are 18.3 kbp or longer).

### Genome assembly

We generated two initial assemblies, one with Falcon and the other with Canu (v1.6)(Koren et al. 2017). The primary Falcon assembly was merged with the canu assembly using quickmerge (Chakraborty et al. 2016), wherein the Canu assembly served as the query. Falcon is a diploid-aware assembler so it can assemble through heterozygous genomic regions that are recalcitrant to Canu. Thus, gaps in the Canu assembly were filled by sequences from the Falcon assembly. This assembly was polished twice with Arrow from SMRT Analysis (v5.1.0) (Chin et al. 2013) and then twice with Pilon (Walker et al. 2014) using 1,203 million 150 bp PE reads (Table S4). The presence of two haplotypes in the raw data may cause the polished assembly to generate redundant sequences if contigs representing alternate haplotypes (i.e. haplotigs) are not identified. To identify alternate haplotigs, we aligned the assembly to itself using Nucmer (-- maxmatch --no-simplify) (Marçais et al. 2018) and identified contigs that were completely embedded within bigger contigs. The sequences in the resulting assembly were marked as either “alt_hap” or “primary” based on whether they were embedded in another contig or not, respectively. While this approach can potentially be confounded by incorrect assembly of repetitive sequences (Phillippy et al. 2008) and aggressively purging alternative haplotigs may remove real duplicate mutations, such adverse outcomes in high quality long-read-based assembly like the *H. charithonia* assembly reported here are rare relative to misassemblies that generate contigs with redundant sequence information (Roach et al. 2018; Guan et al. 2020; Solares et al. 2021). Even so, the placement of rare redundant contigs representing real duplicates is uncertain, diminishing the value in retaining them.

### Microbial decontamination

To decontaminate the microbial sequences from the polished contigs, taxonomic groups were assigned to each contig using Kraken2 (Wood et al. 2019). We identified 4 contigs that were derived from non-butterfly sources (three bacterial and one from nematode). We removed these sequences from the assembly prior to scaffolding and downstream analysis.

### Scaffolding

Hi-C libraries were constructed from a *H. charithonia* female adult whole body. The library was sequenced with PE 75 bp reads on Illumina HiSeq 2500, generating 132,937,739 reads. The reads were mapped to the primary polished and decontaminated assembly using Juicer (Durand et al. 2016) with the default parameters. The contact density map was created from the alignment using the Juicer pipeline and the primary contigs were scaffolded using the Hi-C interaction map following the 3D-DNA pipeline. Among the 70 contigs identified as putatively W-linked (see below), 60 contigs showed Hi-C contacts between them and were joined in a scaffold in Juicebox, following the order suggested by 3D-DNA (Dudchenko et al. 2017). The final assembly contained 21 major scaffolds representing the 19 autosomes, a Z chromosome, and a W chromosome.

### Automated gene annotation

We generated RNA-seq reads from mRNA extracted from antennae, mouthparts and legs of adult *H. charithonia* males and females. Together with previously published RNA-seq data from heads (Catalan et al. 2019), we aligned the reads to the assembly using Hisat2 (Kim et al. 2019). The transcripts were annotated and merged using StringTie (Pertea et al. 2016). We first ran Braker2 (Brůna et al. 2021) to generate a draft annotation based on the *H. charithonia* RNA-seq evidence and protein sequences from *H. melpomene melpomene*. The *H. charithonia* Braker2 annotation, the *H. melpomene* protein and mRNA sequences (Davey et al. 2016), and the *H. charithonia* merged stringtie transcript sequences were used as evidence in Maker2 for gene model prediction (Holt and Yandell 2011). The consensus repeat sequences from Repeatmodeler (see below) was used as the repeat library in Maker2. Maker2 was run in three rounds: in the first run annotation was performed using EST and protein hints, in the second run, Augustus and SNAP predictions were added, and in the third step Genemark predictions were added. The Augustus training was performed in Braker2 and the SNAP prediction was performed using the gene models from the first run of Maker.

### Manual gene annotation

Custom BLAST databases of *H. charithonia* mRNA transcripts were generated from *de novo* (Trinity) and genome-guided transcriptome assemblies of eye, brain, antennae, mouthparts and leg RNA-Seq from adult butterflies. Amino acid sequences for chemosensory proteins (CSPs), odorant binding proteins (OBPs) and olfactory receptors (ORs) identified in *Heliconius* Genome Consortium et al. (*Heliconius* Genome Consortium 2012) and Briscoe et al. (Briscoe et al. 2013) were used as tBLASTn query sequences against this transcriptome in order to identify *H. charithonia* orthologs. Curated OBP, CSP, and OR protein sequences were aligned in MEGA X using MUSCLE. These alignments were visually inspected and manually adjusted. Maximum likelihood trees were estimated in PhyML (Guindon et al. 2010) from the nucleotides using 500 bootstrap replicates and the best-fit substitution models as identified by SMS (Lefort et al. 2017). The Akaike Information Criterion was used as the selection criterion.

### Repeat annotation

We created a custom repeat library using EDTA (Ou et al. 2019) and Repeatmodeler (Flynn et al. 2020). LTR retrotransposons and DNA elements were detected using the EDTA pipeline because EDTA is more accurate at finding intact elements than Repeatmodeler. In EDTA, we used the *H. charithonia* protein sequences from the final Maker run for filtering out predicted TEs that overlapped protein coding sequences. Because EDTA does not annotate non-LTR retrotransposons, the non-LTR elements were identified using Repeatmodeler and added to the repeat library.

### Identification of W-linked sequences

To identify the W-linked sequences, male and female Illumina paired-end genomic DNA reads were aligned to the polished and decontaminated contig assembly using Bowtie2 (v2.2.7)(Langmead and Salzberg 2012). Alignments were sorted and male and female Illumina read coverage (Table S4) of each contig was measured using Bedtools (bedtools coverage - mean)(Quinlan and Hall 2010) and contigs showing at least 2-fold higher coverage for female reads than male reads were designated as putative W-linked contigs. The contigs showing >2 fold male-to-female coverage ratio were assigned as the candidate Z contigs. This Z chromosome candidate mapped to the *H. erato* Z chromosome, suggesting that the coverage based sex-chromosome assignment identified sex-linked chromosomes correctly (Fig. S4). Contigs showing enrichment of female k-mers were marked as candidates for W-linked sequences. Finally, we mapped the RNA-seq reads from males and females to repeat-masked putative W-linked sequences and compared the male vs. female transcript abundance in the putative W-linked genes.

### *UVRh1* PCR amplification

To examine the sex-linkage of *UVRh1* in 10 *Heliconius* species, genomic DNA was extracted from the dissected thorax of single adult male and female butterflies from each species using Monarch Genomic DNA Purification Kit (New England Biolabs) following the manufacturer’s protocol, except we added 10 uL of proteinase K to each sample. To amplify *UVRh1* genomic sequence, we used the primer pairs 5’ CGCTACAGTCTTGCAAGCTAC 3’ and 5’ ATATTTCTACAGTGGAATCGTAAAA 3’. For all amplifications using the *UVRh1*-specific primers, we used Phusion HF Polymerase (New England Biolabs) and annealing temperatures (Tm) of 60°C and 58°C, respectively. To rule out missing amplicons due to PCR failure in the fresh genomic DNA samples, we used the forward primer (ef44) 5’ GCYGARCGYGARCGTGGTATYAC 3’ and reverse primer (efrcM4) 5’ ACAGCVACKGTYTGYCTCATRTC 3’ to amplify the housekeeping gene *EF1a*. The purified *UVRh1* amplicons were cloned into the minT vector using the PCR cloning kit and following the manufacturer’s protocol (NEB). The cloned amplicons were sequenced by Retrogen Inc. using the NEB-F, NEB-R primers supplied by the manufacturer.

### Ancestral state reconstruction

Presence or absence of *UVRh1* mRNA or protein expression in adult male *Heliconius* eyes was determined based on RNA-seq data of McCulloch et al. (2017), reproduced in Table S1 and and/or immunohistochemistry shown in Fig. S7. Characters were mapped on a trimmed *Heliconius* species phylogeny (Kozak et al. 2015) using Mesquite v.3.10. Ancestral state likelihood analysis was performed in Mesquite using binary character states.

### *UVRh1* knockout using CRISPR

To knock out *UVRh1* using CRISPR (Jinek et al. 2012), we designed two gRNAs (5’ GGAGTACAGCAACGCTAGTG 3’, 5’ GGTTTTGCTACAGGTGCTTT 3’) that target the second and third exons of *UVRh1*, respectively. The gRNAs were synthesized (Synthego) and were combined with Cas9 (EnGen® Spy Cas9 NLS, New England Biolabs) at concentrations 160 ng/uL and 240 ng/uL, respectively.

Embryos were collected by giving fresh young *Passiflora biflora* shoots to adults for one hour and the collected embryos were soaked in 5% benzylkonium chloride solution (Millipore Sigma) for 5 minutes for disinfection. The gRNA-Cas9 mixture was incubated at room temperature for 10 minutes for formation of ribonucleoprotein complex and was injected into 0-1.5h embryos attached to a double-sided tape on a glass slide. Injected embryos were kept inside a petri dish for 4 days at room temperature with moistened Kimwipes to maintain humidity. Eggs hatched after ∼4 days and the ∼4 days old caterpillars were transferred to a *P. biflora* inside a mesh cage. Adults eclosed after approximately four weeks and were genotyped for the CRISPR mediated deletion using PCR.

To screen adults for the CRISPR-mediated deletion, we extracted genomic DNA from a hind leg of each adult using Monarch Genomic DNA Purification Kit (New England Biolabs) and amplified the DNA using a *UVRh1*-specific primer pair (5’ CAAGCATTTGTCATTGATGCA 3’, 5’ GAAACGCAAAACTACAACGTT 3’) that produced a 708 bp and 390 bp amplicons for uncut and cut *UVRh1* genomic sequences, respectively.

### Immunohistochemistry of adult eyes

Methods were adapted from previous studies (Hsiao et al. 2012; Perry et al. 2016; McCulloch et al. 2017). Dissected *H. charithonia* eyes were fixed in 4% paraformaldehyde (in 1x PBS) for one hour at room temperature with one hour baths at room temperature in increasing concentrations of sucrose (10, 20, and 30%) afterwards. The corneal lens was then excised from each eye and the eyes were embedded in blocks of gelatin-albumin. The blocks were then fixed in 4% formalin (in 1x PBS) for six hours and a VF-310-0Z Compresstome (Precisionary) was used to cut 50 μm slices. Tissue slices were blocked for one hour in 10% (v/v) normal goat serum and normal donkey serum and 0.3% Triton X-100 (in 1X PBS). Tissues were incubated with preadsorped primary antibodies (1:15 guinea pig anti-UVRh1, 2:75 rabbit anti-UVRh2, and 1:15 chicken anti-BRh in blocking solution) overnight at 4°C. Tissues were washed 5X 15 minutes in 1x PBS and incubated overnight at 4°C with secondary antibodies (1:250 goat anti-guinea pig AlexaFluor 633, 1:250 donkey anti-rabbit Cy3, and 1:250 goat anti-chicken AlexaFluor 488 in blocking solution). Afterwards, tissues were then washed 5x 15 minutes in PBST and then mounted in 70% glycerol. Images were taken using a Zeiss LSM 900 Airyscan 2 confocal microscope under a 20x/0.8NA dry objective in the UC Irvine Optical Core Facility, exported using ZenBlue 3.5, and processed/pseudocolored using Fiji (Schindelin et al. 2012).

### Behavioral trials

Both 390 nm and 380 nm 10 nm bandpass filtered lights were on during training at 1:1 intensity but only 390 nm light was rewarded with 10% honey water supplemented with pollen (+) while the unrewarded light had water (-). After training, both sexes (n=3 individual butterflies per sex) were then tested for UV discrimination ability between 390 nm (+) and 380 nm (−) over three different intensity combinations where the relative intensity of the rewarded:unrewarded lights was 1:5, 1:1, or 5:1 (n=15 trials per intensity). During training and between training sessions, the placement of the rewarded and unrewarded stimuli was randomly switched so that the butterfly did not learn to associate the position of the light with a reward. The apparatus was cleaned after each session with 70% isopropyl alcohol to remove chemical cues. After about 4– 5 days of training, butterflies were capable of independently flying toward the apparatus and making a choice between the two light stimuli. Three different approximate ratios of the peak physical intensities or absolute brightnesses of the rewarded/unrewarded stimuli were used: 0.02, 1.0 and 5.0 (or 1:5, 1:1, and 5:1)(Fig. S10). Butterflies first completed trials at an intensity combination of 1:1 (15 choices each). Following this test they were given random choices between intensities of 1:5 or 5:1 (rewarded:unrewarded) until they had completed 15 choices with each intensity combination. The number of correct versus incorrect choices each butterfly made at different intensity combinations was modeled using a general linear model with Poisson distribution in R statistical software (version 4.1.1).

