## Supplementary Material for "Sex-linked gene traffic underlies the acquisition of sexually dimorphic UV color vision in *Heliconius* butterflies"

### Supplementary text and figures

#### Manual annotation of transcriptome

##### Gene discovery, curation, and nomenclature

Manual annotations of chemosensory-related proteins were conducted in part to assess the completeness of the genome assembly. Protein sequences of known *Heliconius melpomene* chemosensory proteins (CSPs), odorant binding proteins (OBPs), and olfactory receptors (ORs) [1] were queried against both *de novo* (HCH_heads_denovo.fasta, HCH708_antenna_trinity.fasta) and genome-guided (Stringtie.fasta) *Heliconius charithonia* RNA-seq transcriptome assemblies deposited in Dryad (https://doi.org/XX.XXXX/dryad.XXXX) using the tBLASTn search algorithm.

Blast results, sorted by ascending E-value and using the default E-value cutoff, were visually inspected. Top hits were imported, trimmed, and aligned in MEGA X (Kumar et al. 2018). *H. charithonia* genes were numbered according to their closest *H. melpomene* homologs following phylogenetic analysis (see below). Alternate blast hits subjectively determined to be of significant length and similarity were also added to the alignments to explore potential expansions of these gene families. All annotated transcripts were deposited in GenBank under accession nos: XXXXXXXX-XXXXXXXX.

##### Phylogenetic Analysis

Curated OBP, CSP, and OR protein sequences from *H. melpomene* and *H. charithonia* were aligned in MEGA X using the MUSCLE algorithm. These alignments were visually inspected and manually adjusted as needed. Maximum likelihood trees were estimated in PhyML (Guindon et al. 2010) from the nucleotides using 500 bootstrap replicates and the best-fit substitution models as identified by SMS (Lefort, Longueville, and Gascuel 2017). The Akaike Information Criterion was used as the selection criterion. Tree images were generated using the iTOL web server (Letunic and Bork 2019).

##### Genome Analysis

We also searched the *H. charithonia* genome for the same CSPs, OBPs, and ORs (some of which were extracted directly from the genome). For those genes where we found an *H. charithonia* ortholog transcript, we used the ortholog's nucleotide sequence as a query for a BLASTn search in the genome. Where there was no ortholog found, we searched using tBLASTn with the *H. melpomene* protein sequence. Blast results were visually inspected, and relevant hits were determined in the same way as described for the transcriptome searches.

**Chemosensory proteins (CSPs)**

Out of the 33 *H. melpomene* CSPs reported in *Heliconius* Genome Consortium (*Heliconius* Genome Consortium 2012), we identified 28 orthologs in *H. charithonia (Hc)*. 23 of these orthologs are full-length, containing a start and stop codon. We also identified 8 lineage-specific CSPs, 7 of which are full-length. This makes 36 *Hc* CSPs identified in total. In the *Hc* genome, we found genes corresponding to individual mRNAs for all of the 28 ortholog transcripts, with 20 of these genes being full-length. We also located in the genome all of the 8 new *Hc* CSPs, with 7 of these being full-length.

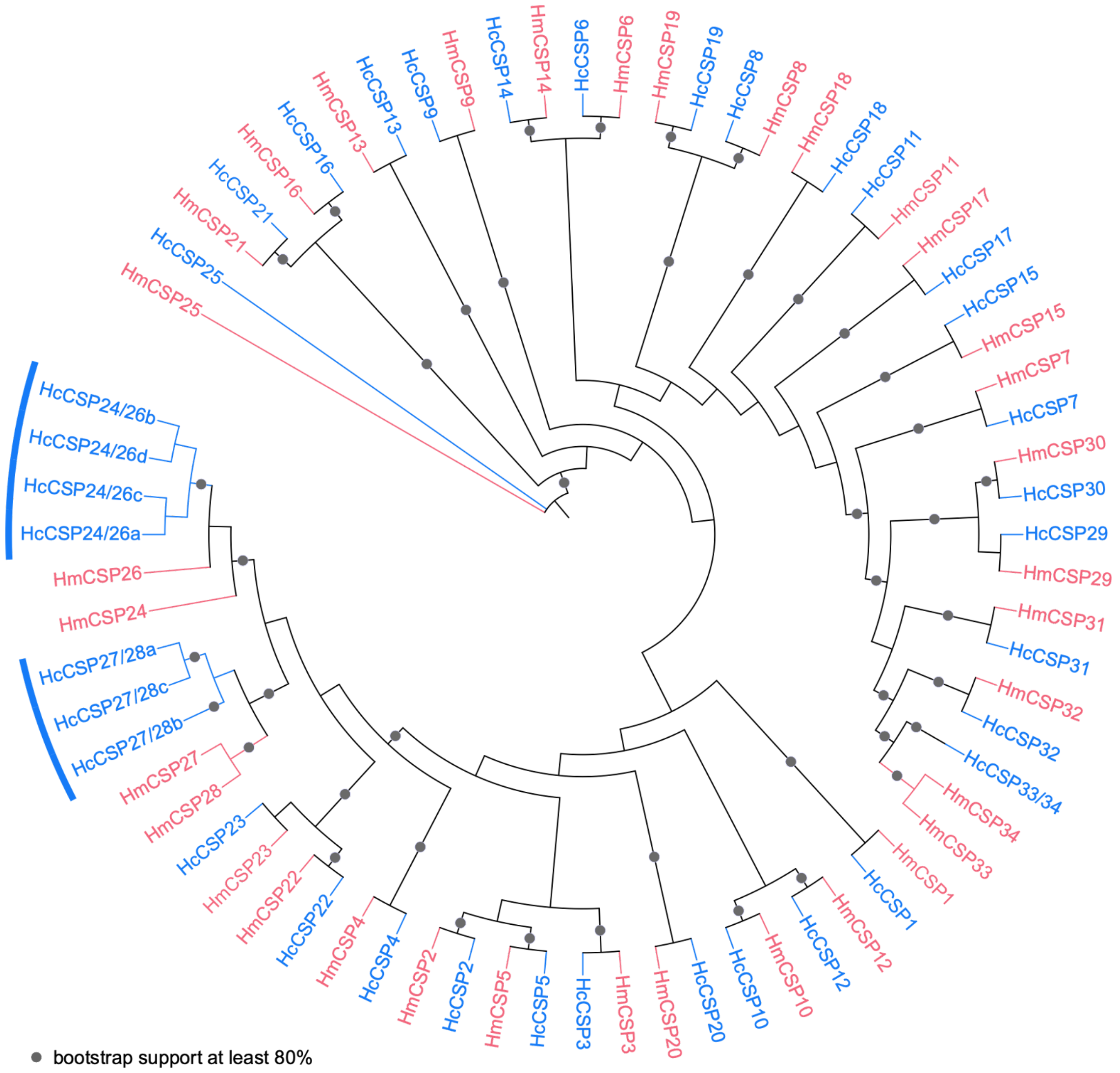

**Fig. S1. Maximum likelihood tree of manually annotated chemosensory proteins (CSPs).** Genes from *Heliconius melpomene* -and *H. charithonia* are shown in red and blue, respectively. Amino acid sequences were aligned, then backtranslated to nucleotides to build the tree. Grey circles on branches indicate bootstrap values >= 80% from 500 bootstrap replicates. Branches highlighted by red or blue arcs indicate lineage-specific CSP expansions. The model selected for phylogenetic analysis was GTR+G. Hm, *Heliconius melpomene*; Hc, *Heliconius charithonia*.

**Odorant-binding proteins**

While 43 putative *H. melpomene* OBPs were reported in *Heliconius* Genome Consortium (*Heliconius* Genome Consortium 2012), mRNA nucleotide transcripts were only available for 41 of them. Using these 41 known OBPs as reference (*Heliconius* Genome Consortium 2012), we identified 35 *H. charithonia* OBP orthologs. 26 of these orthologs were full-length, containing a start and stop codon.

We also identified 4 lineage specific *H. charithonia* OBPs, all of which were full-length. In total, this makes 39 *H. charithonia* OBPs. In the *Hch* genome, we found genes corresponding to individual mRNAs for all 35 ortholog transcripts, and for each of the 4 new OBP transcripts.

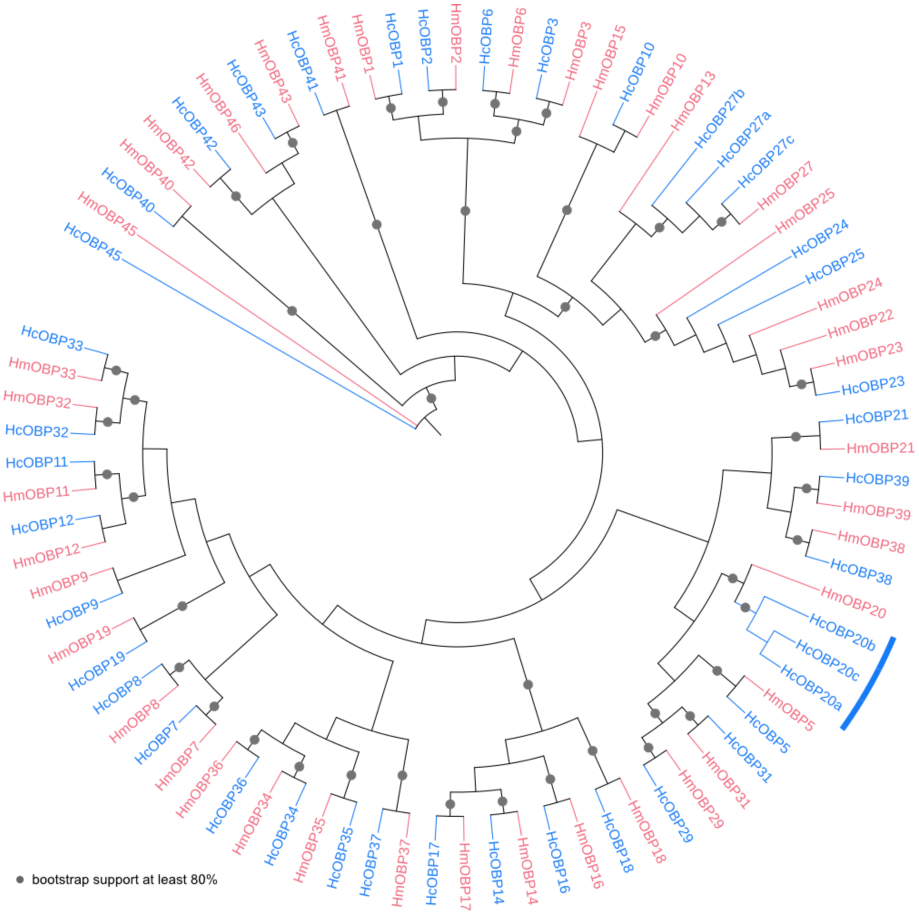

**Fig. S2. Maximum likelihood tree of manually annotated odorant-binding proteins (OBPs).** Genes from *Heliconius melpomene* -and *H. charithonia* are shown in red and blue, respectively. Amino acid sequences were aligned, then backtranslated to nucleotides to build the tree. Grey circles on branches indicate bootstrap values >= 80% from 500 bootstrap replicates. The model selected for phylogenetic analysis was GTR+G+I. Hm, *Heliconius melpomene*; Hc, *Heliconius charithonia*.

#### Olfactory receptors

While 70 putative *H. melpomene* OR genes were reported in *Heliconius* Genome Consortium (*Heliconius* Genome Consortium 2012), mRNA nucleotide transcripts were only available for 52 of them. We identified orthologs in *H. charithonia* for 50 of these 52 ORs. Of the 50 orthologs, 11 were full-length, containing a start and stop codon. We also found 5 lineage-specific ORs, 1 of which was full-length. Altogether this makes 55 *H. charithonia* ORs. In the *Hc* genome, we found genes corresponding to the individual mRNAs for all 55 ORs. One previously identified OR in *H. melpomene, HmOR65,* is actually identical to *HmOR51*. A BLASTp against GenBank for this sequence yielded only *HmOR51*. For that reason, we omitted *HmOR65* from our analysis.

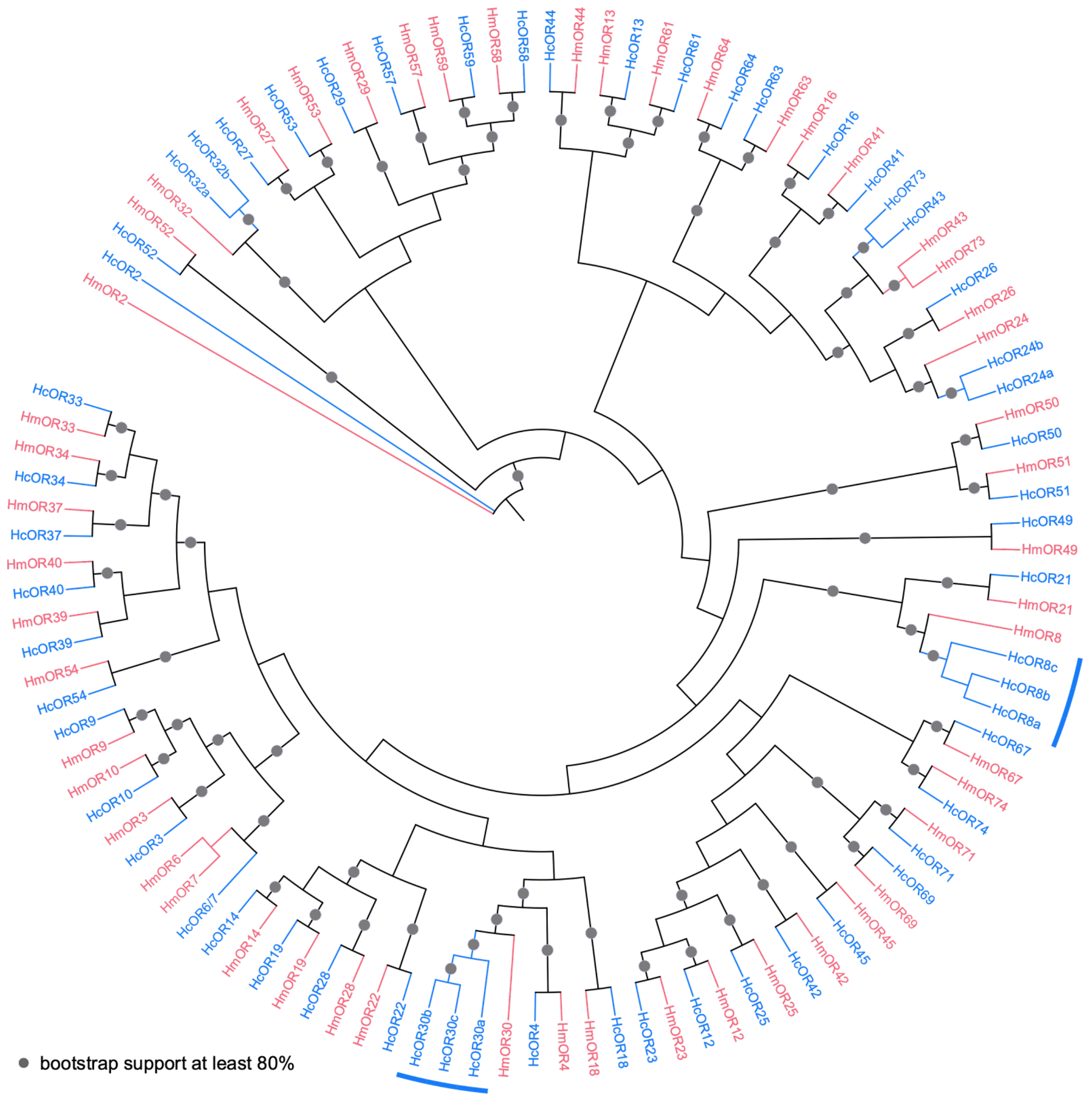

**Fig. S3. Maximum likelihood tree of manually annotated olfactory receptor proteins (ORs).** Olfactory receptors from *Heliconius melpomene* and *H. charithonia* are shown in red and blue, respectively. Amino acid sequences were aligned, then backtranslated to nucleotides to build the tree. Grey circles on branches indicate bootstrap values >= 80% from 500 bootstrap replicates. The model selected for phylogenetic analysis was GTR+G. Hm, *Heliconius melpomene*; Hc, *Heliconius charithonia*.

A

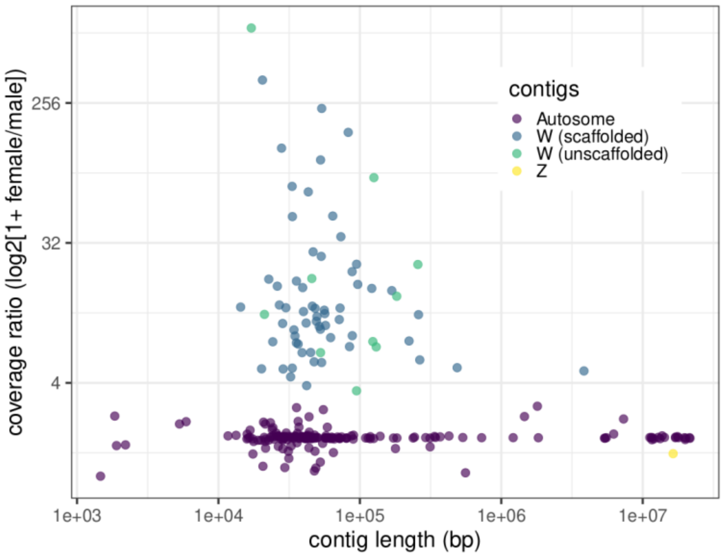

B

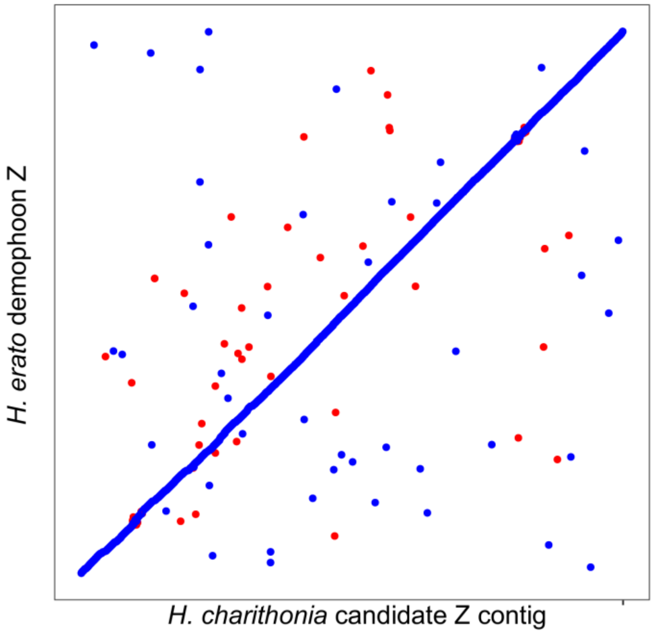

**Fig. S4. Identification of Z contig.** A) The contig showing >2 fold male-to-female coverage ratio was assigned as the candidate Z contig. B) Alignment dot plot between the *H. charithonia* Z chromosome candidate contig and *H. erato demophoon* Z chromosome scaffold (Nadeau et al. 2014). Mapping of the Z chromosome candidate to the *H. erato* Z chromosome suggests that the coverage based sex-chromosome assignment identified sex-linked chromosomes correctly.

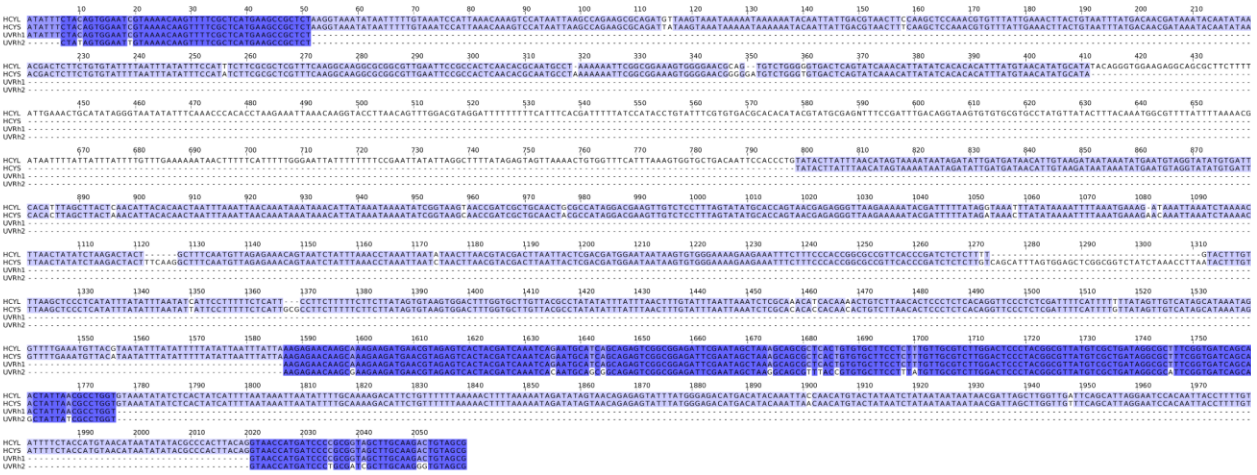

**Fig. S5.** Muscle alignment of long (HCYL) and short (HCYS) *UVRh1* amplicons from a female *H. cydno* (Fig. 4B) and corresponding exonic sequences from *UVRh1* (GenBank ID GQ451895.1) and *UVRh2* (Genbank ID GQ451896.1) cDNA from the same species. As evident from shared homology between the amplicons and *UVRh1*, both amplicons are from *UVRh1*. As the longer *UVRh1* amplicon (HCYL) is present only in the female studied here, *H. cydno* either carries a female-specific *UVRh1* or the two autosomal *UVRh1* alleles in the female are segregating for an indel structural variant.

A

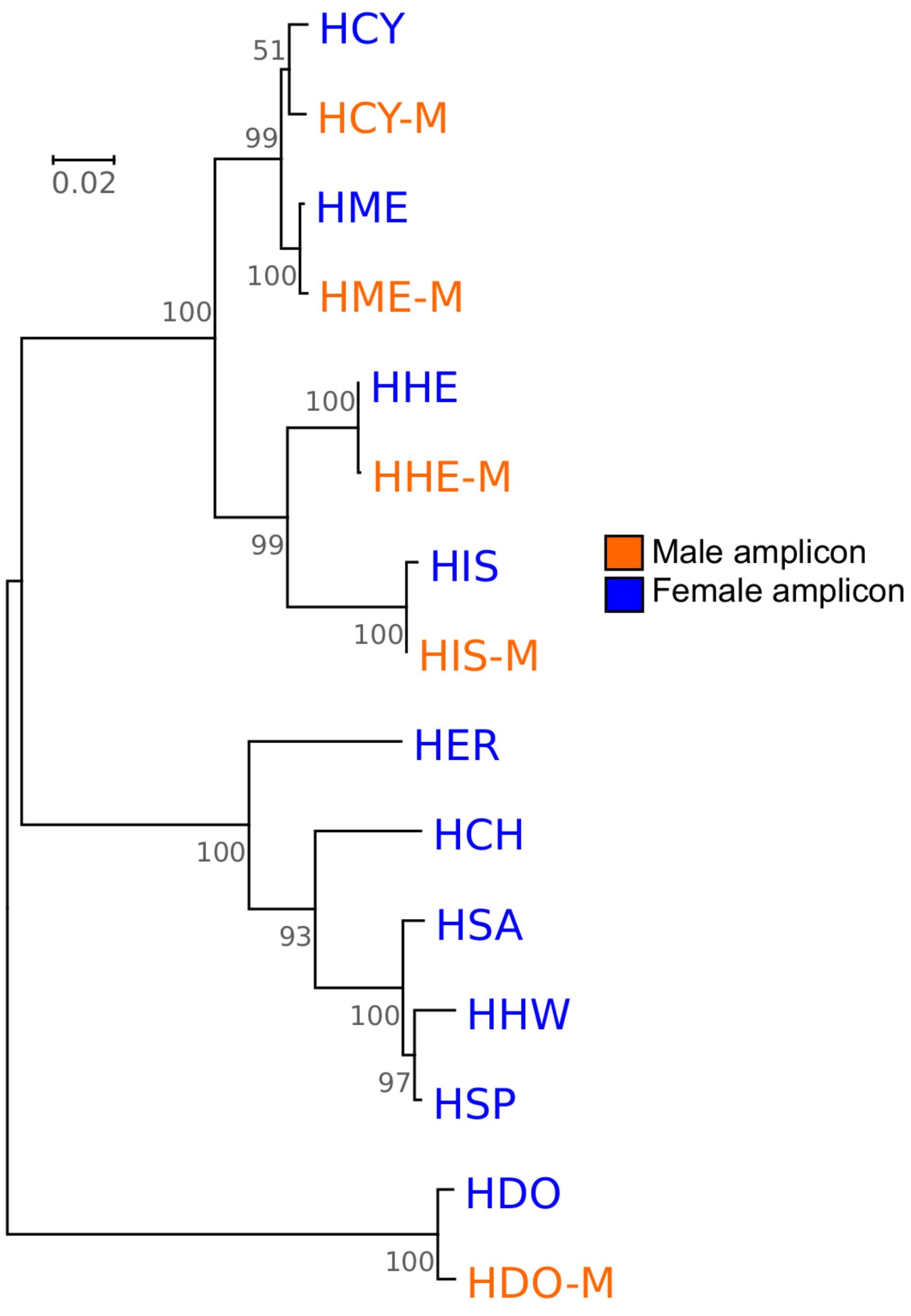

0.02
HCY
HCY-M
HME
HME-M
HHE
HHE-M
HIS
HIS-M
HER
HCH
HSA
HHW
HSP
HDO
HDO-M
51
99
100
100
100
99
100
100
93
100
97
100
Male amplicon
Female amplicon

B

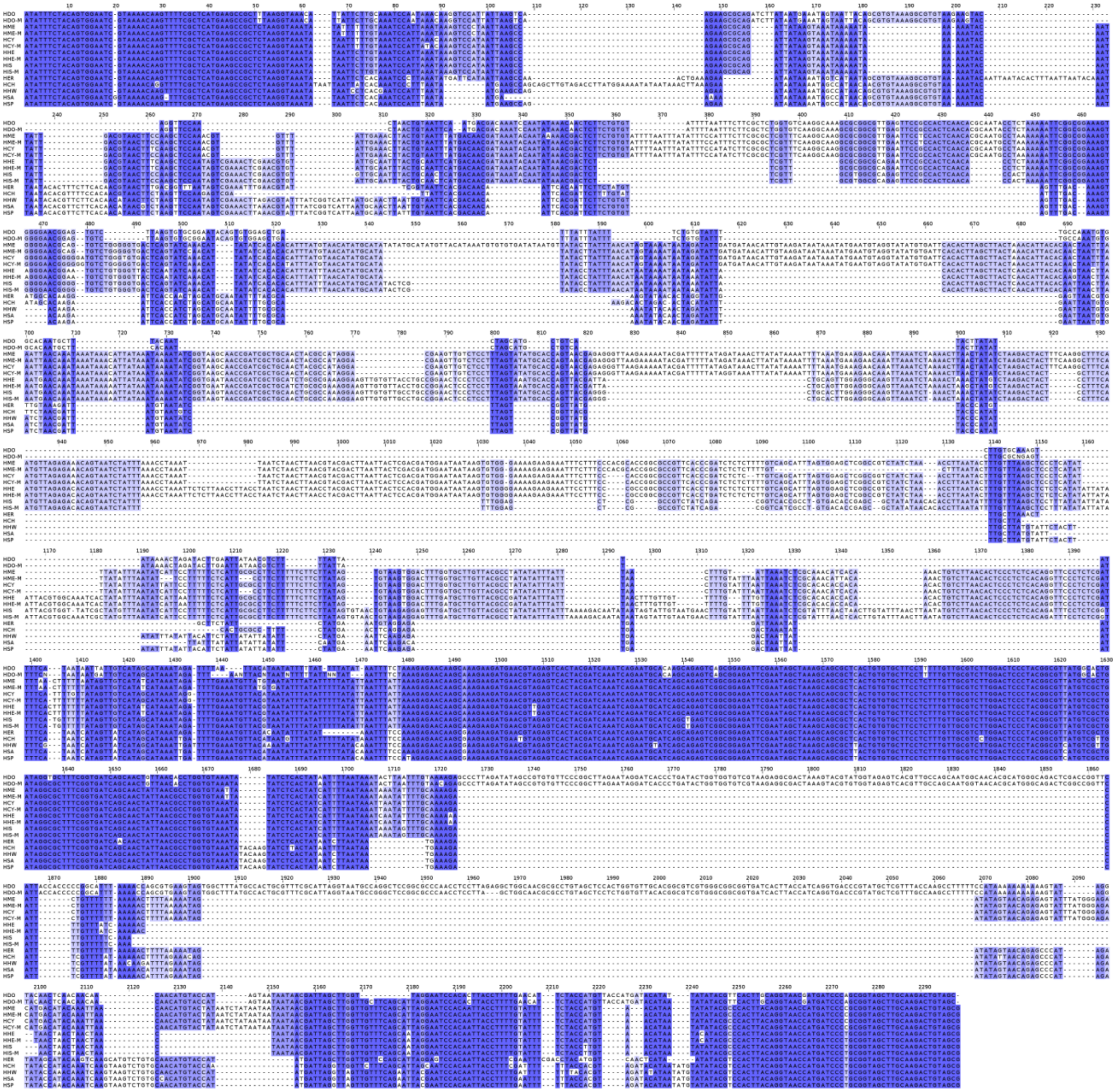

**Fig. S6. Phylogeny of sequenced PCR products amplified from gDNA of various *Heliconius* species using *UVRh1* primers.** A) Phylogenetic relationship of *UVRh1* sequences shown in Fig. 3 based on ML analysis of 2294 positions. The tree with the highest log likelihood (-6146.44) is shown. The tree is drawn to scale, with branch lengths measured in the number of substitutions per site. B) Alignment of 2294 nucleotide positions of *UVRh1* used in the phylogenetic analysis. Abbreviations: HCH, *H. charithonia*; HCY, *H. cydno;* HDO, *H. doris*, HER, *H. erato*; HHE, *H. hecale*; HHW, *H. hewitsoni*; HME, *H. melpomene*; HSA, *H. sara;* HSP, *H. sapho*. B) Muscle alignment of sequenced *UVRh1* amplicons.

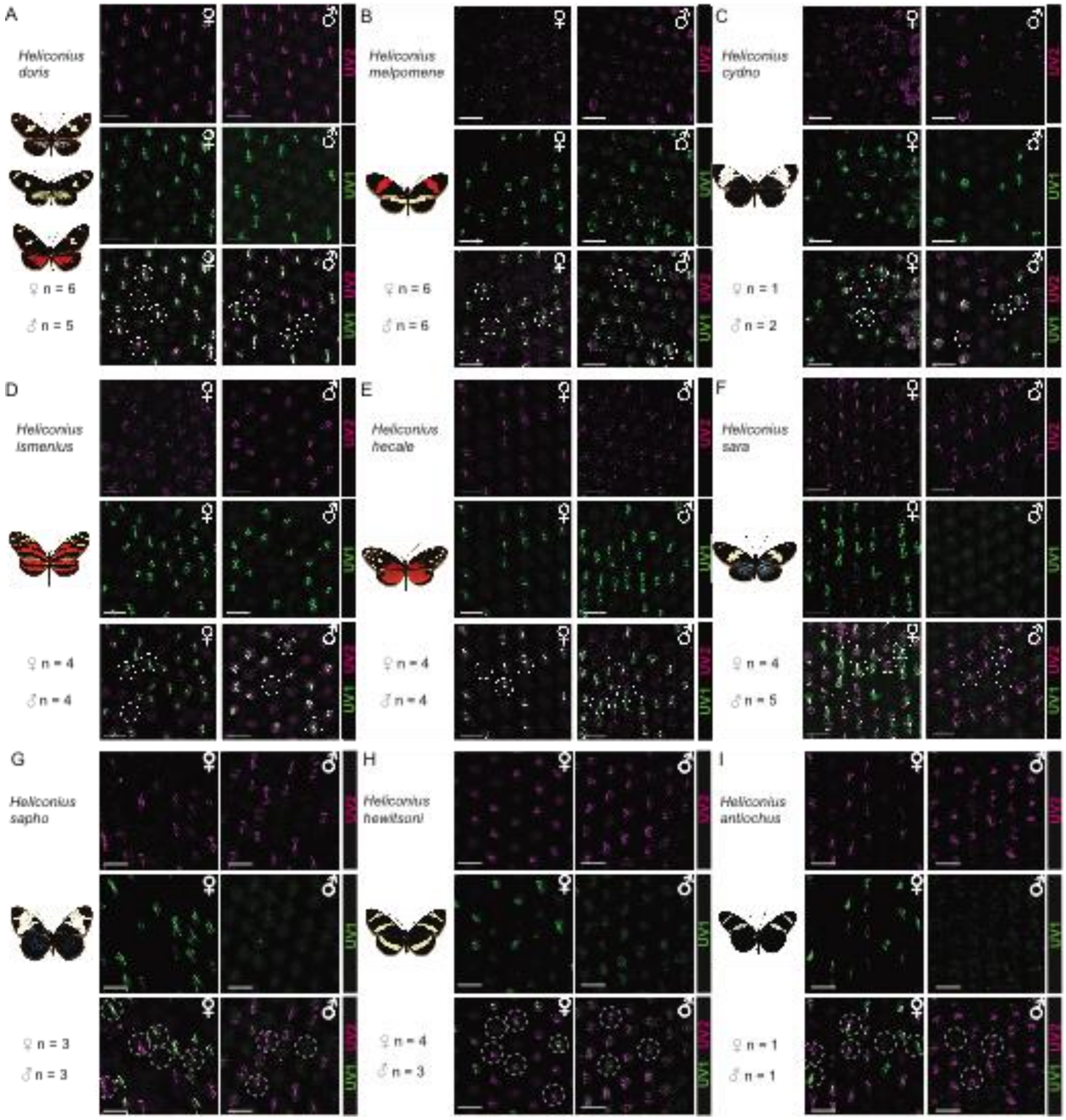
A
Heliconius doris
♀ n = 6
♂ n = 5
B
Heliconius melpomene
♀ n = 6
♂ n = 6
C
Heliconius cydno
♀ n = 1
♂ n = 2
D
Heliconius ismenius
♀ n = 4
♂ n = 4
E
Heliconius hecale
♀ n = 4
♂ n = 4
F
Heliconius sara
♀ n = 4
♂ n = 5
G
Heliconius sapho
♀ n = 3
♂ n = 3
H
Heliconius hewitsoni
♀ n = 4
♂ n = 3
I
Heliconius antiochus
♀ n = 1
♂ n = 1
UV2
UV1

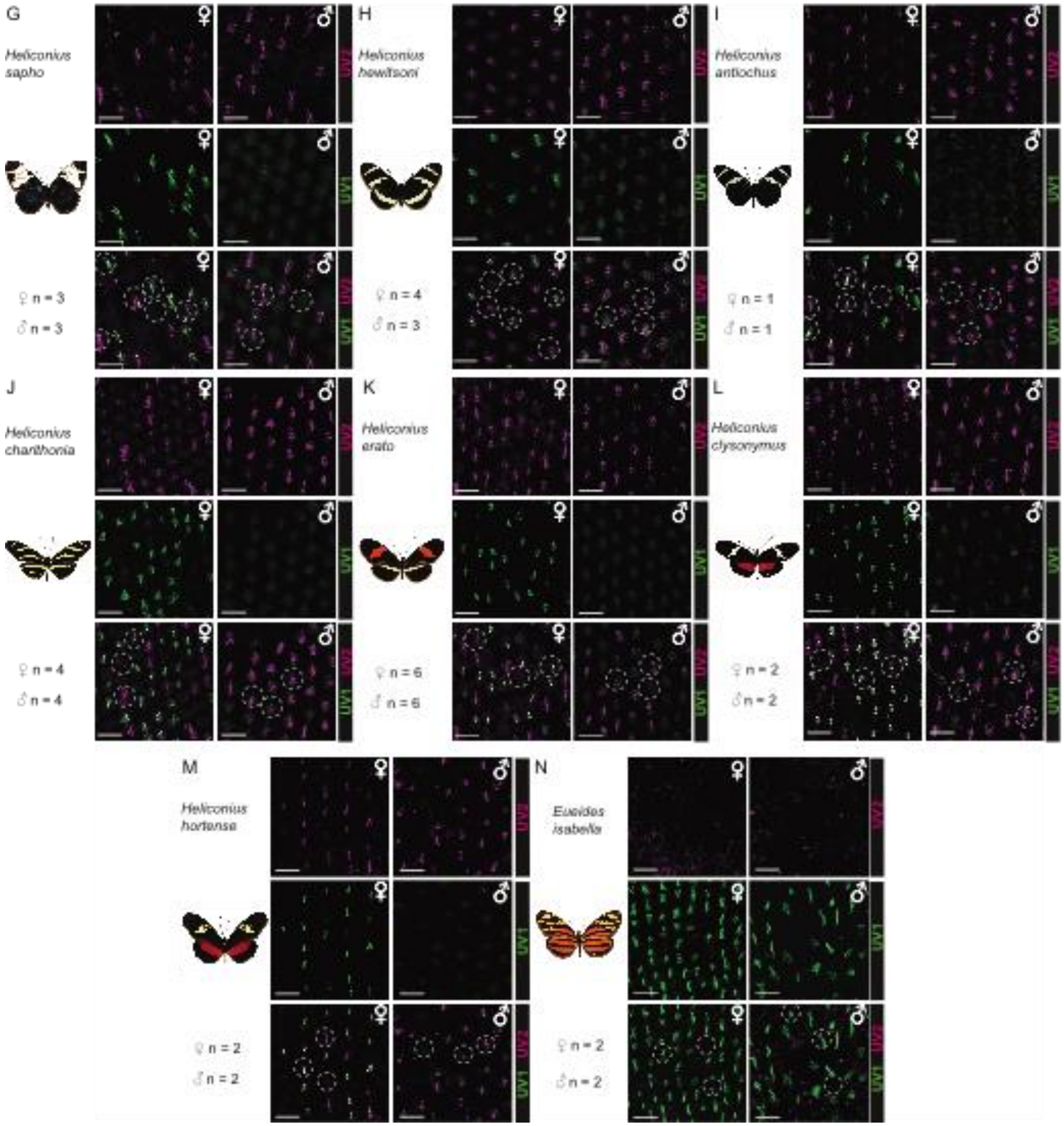

**Fig. S7. Eye sections of adult *Heliconius* and outgroup species immunostained for UVRh1 (green) and UVRh2 (magenta) opsins**. (A-N) Images of each sex for all immunostained species. Sample size refers to number of individuals sampled per species and sex. Dashed circles identify the different types of ommatidia in each species and sex. Scale bars, 25 µm. Methods used to produce these images are described in McCulloch et al. 2017.

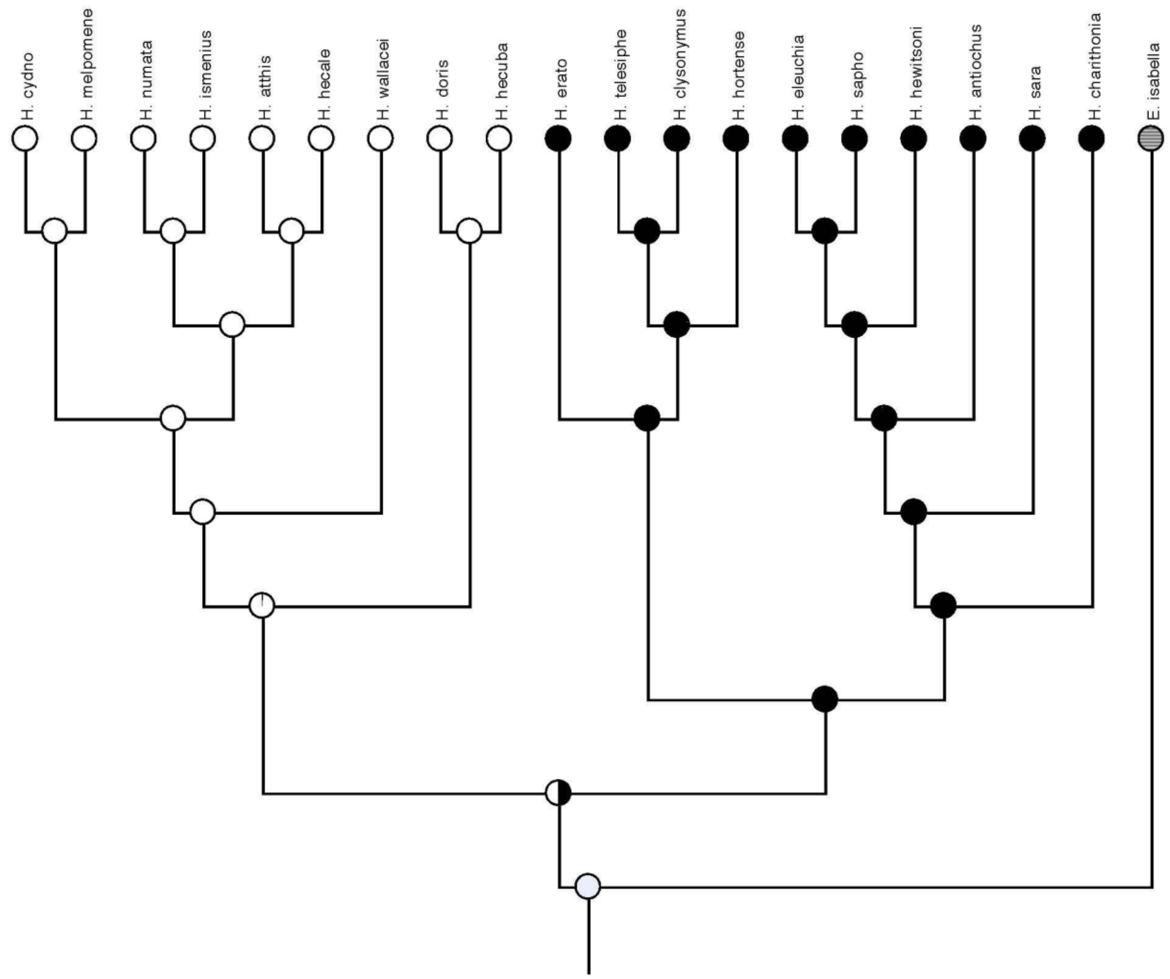

**Fig. S8. Ancestral state reconstruction of presence or absence of UVRh1 mRNA/protein expression in males.** Presence (black) or absence (white) of male *UVRh1* mRNA or protein expression was determined via the analysis of RNA-seq from eye+brain tissue in Table S1 and/or IHC of adult eye tissue using anti-UVRh1 and anti-UVRh2 antibodies shown in Fig. S7 (see also McCulloch et al. 2017). Species where the average *UVRh1* reads per kilobase per million (RPKM) for males was <1, were scored as absent. For species in which *UVRh1* RPKM >1, then *UVRh1* mRNA was scored as present in males.

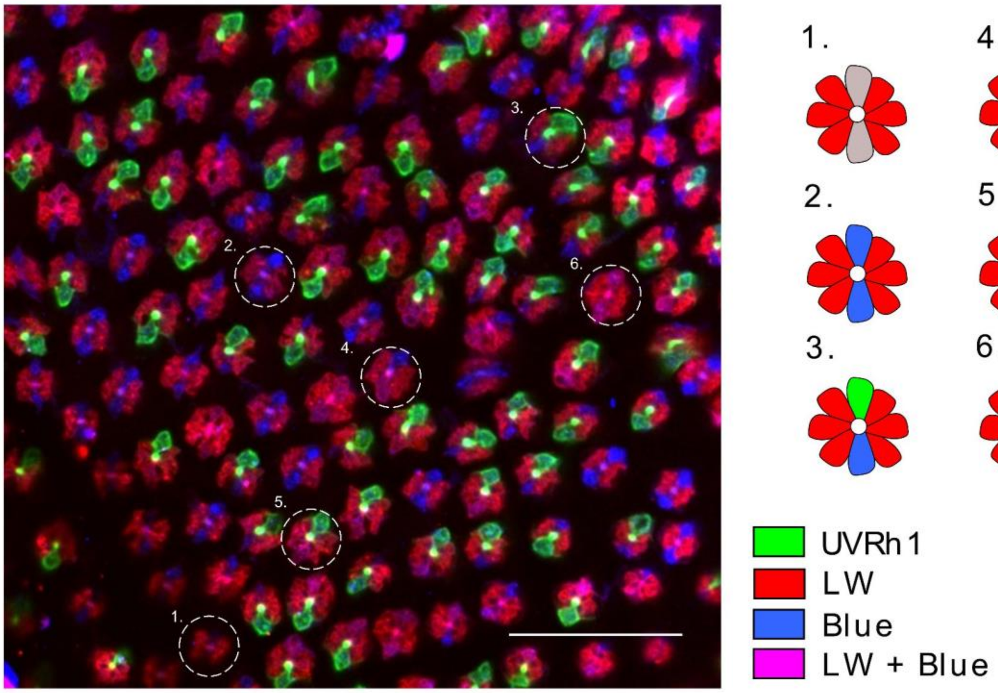

**Fig. S9. UVRh1, BRh and LWRh opsin expression in a wild-type adult female *H. charithonia* compound eye.** (Right) Anti-UVRh1 (green), anti-LWRh (red) and anti-BRh (blue) antibody staining of a female retina. (Left) Adult *H. charithonia* female retinas have at least six types of ommatidia based on opsin expression in the R1 and R2 photoreceptor cells shown here and in Fig. 4D, including ommatidial types which co-express LWRh and BRh: 1. UVRh2/UVRh2, 2. BRh/BRh, 3. UVRh1/BRh, 4. BRh/LWRh-BRh, 5. UVRh1/LWRh-BRh, 6. LWRh-BRh/LWRh-BRh. For evidence of LWRh-BRh co-expression in other *Heliconius* species see (McCulloch et al. 2022). Scale bars, 50 µm.

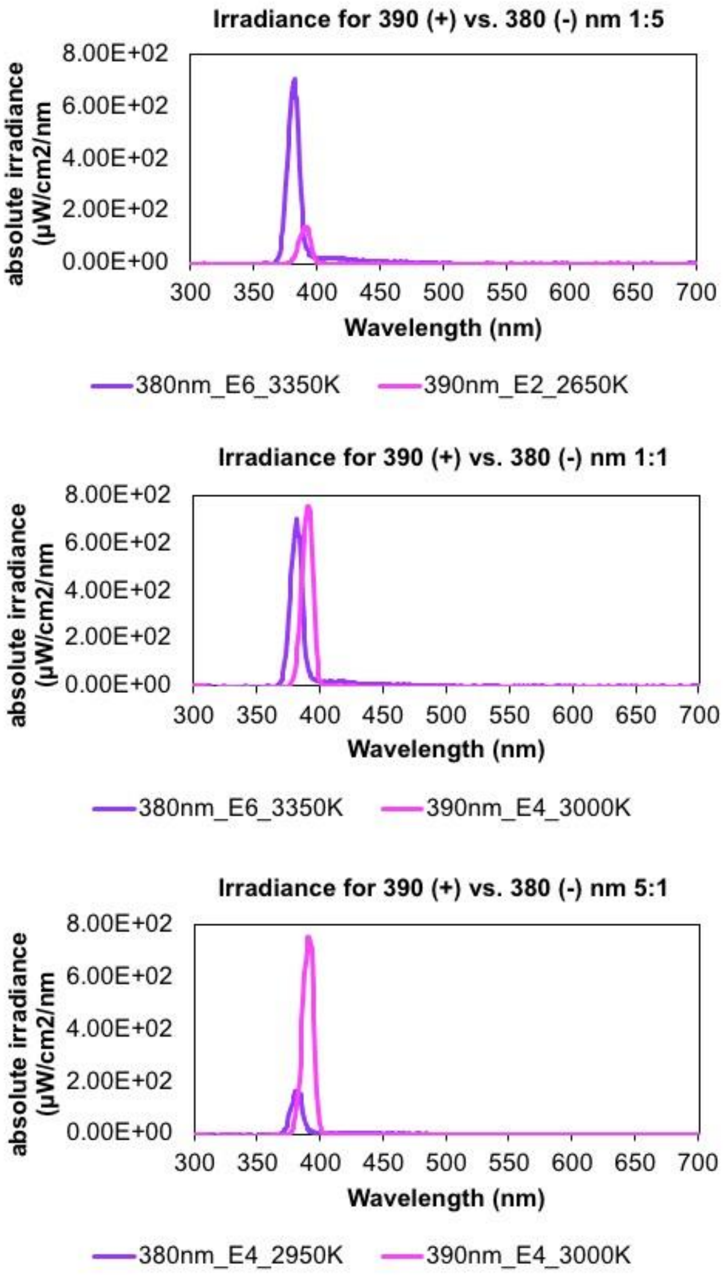

**Fig. S10. Absolute irradiance spectra for 390 nm and 380 nm filtered lights used for butterfly behavior training and testing.** Lights measured using an Ocean Optics USB2000 spectrometer and 100 µm-diameter fiber optic cable. (A) Irradiance for training or testing 390 nm (+) vs. 380 (-) nm 1:1. (B) Irradiance for testing 390 nm (+) vs. 380 nm (-) 1:5. (C) Irradiance for testing 390 nm (+) vs. (380 nm) (-) 5:1.

#### Supplementary Tables

**Table S1.** Sex-specific differential *UVRh1* expression in *Heliconius* based on RNA-seq read-mapping from McCulloch et al. (2017). Sexually dimorphic *UVRh1* mRNA is defined as being present in species where the average RPKM of UV1 in males is <1.

| | | Males | | | Females | | |
|---|---|---|---|---|---|---|---|
| **Clade** | **Species** | **N** | **Average RPKM UV1** | **Average RPKM UV2** | **N** | **Average RPKM UV1** | **Average RPKM UV2** |
| *doris* | *H. doris* | 3 | 385.67 | 718 | 3 | 693.62 | 651.7 |
| | *H. hecuba* | 1 | 1760.67 | 1014.5 | 1 | 3214.53 | 290.35 |
| *wallacei* | *H. wallacei* | 1 | 979.51 | 921.8 | 0 | ND | ND |
| *melpomene* | *H. melpomene* | 2 | 684.98 | 181.3 | 2 | 554.12 | 66.35 |
| silvaniform | *H. atthis* | 1 | 2114.9 | 11.9 | 1 | 4075.18 | 0.61 |
| | *H. hecale* | 2 | 334.59 | 4.5 | 2 | 1184.19 | 6.07 |
| | *H. ismenius* | 2 | 807.98 | 3.1 | 2 | 971.14 | 1.65 |
| | *H. numata* | 2 | 637.35 | 8.5 | 1 | 757.06 | 6.29 |
| *aoede* | *H. aoede* | 1 | 0 | 4463.7 | 0 | ND | ND |
| *sara* | *H. charithonia* | 2 | 0.77 | 2359.3 | 3 | 748.48 | 1210.36 |
| | *H. eleuchia* | 1 | 0.17 | 4631.06 | 1 | 423.12 | 5515.05 |
| | *H. hewitsoni* | 2 | 0.68 | 1105.2 | 2 | 373.13 | 1327.59 |
| | *H. sapho* | 1 | 0.11 | 1387 | 1 | 277.85 | 1039.18 |
| | *H. sara* | 3 | 0.36 | 1349.6 | 4 | 284.58 | 1152.47 |
| *erato* | *H. clysonymus* | 1 | 0.93 | 3879.8 | 2 | 1043.01 | 1033.01 |
| | *H. erato* | 2 | 0.35 | 1886.1 | 2 | 519.83 | 867.88 |
| | *H. hortense* | 1 | 0.13 | 2664.2 | 1 | 1491.18 | 1507.52 |
| | *H. telesiphe* | 1 | 0.52 | 1515.1 | 1 | 847.3 | 1037 |

Uniquely-mapped reads to each *UVRh* opsin mRNA were quantified by calculating reads per kilobase of transcript per million mapped (RPKM).

**Table S2.** Choice data for rewarded (390 nm)(+) versus unrewarded (380 nm) (-) lights for individual female and male *H. charithonia* butterflies over three light intensity treatments: 1(+): 5(-), 1(+):1(-), and 5(+):1(-). N=3 butterflies/sex/light intensity treatment; 15 choices per butterfly/light intensity treatment=45 choices total per light intensity treatment.

| Species | Sex | Light intensity | Correct choices out of 15 |
|---|---|---|---|
| *charithonia* | F | 1:5 | 11 |
| | F | 1:5 | 10 |
| | F | 1:5 | 11 |
| | F | 1:1 | 9 |
| | F | 1:1 | 12 |
| | F | 1:1 | 11 |
| | F | 5:1 | 12 |
| | F | 5:1 | 11 |
| | F | 5:1 | 9 |
| | M | 1:5 | 1 |
| | M | 1:5 | 4 |
| | M | 1:5 | 5 |
| | M | 1:1 | 3 |
| | M | 1:1 | 6 |
| | M | 1:1 | 9 |
| | M | 5:1 | 10 |

| | | | |
|---|---|---|---|
| | **M** | **5:1** | **12** |
| | **M** | **5:1** | **10** |

**Table S3** A GLM model with poisson distribution as implemented in R v 4.1.1 was used to test the hypothesis that *H. charithonia* individuals significantly chose the rewarded light, 390 nm, over the unrewarded light, 380 nm. Three individuals per sex per rewarded: unrewarded light intensity treatment were given 15 choices each for a total of 45 choices.

| **Species** | **Sex** | **N choices** | **Light Condition** | **Z value** | **p-value** |
|---|---|---|---|---|---|
| *H. charithonia* | F | 45 | 1:5 | 2.739 | 0.01 |
| | F | 45 | 1:1 | 2.739 | 0.01 |
| | F | 45 | 5:1 | 2.739 | 0.01 |
| | M | 45 | 1:5 | -3.494 | 0.001 |
| | M | 45 | 1:1 | -1.332 | 0.18 |
| | M | 45 | 5:1 | 2.739 | 0.01 |

**Table S4**. List of specimens and localities used in DNA- and RNA-seq and PCRs. Tissue type, sequencing type, assembly (if available), accession number are listed.

| Species | Origin | ID | Sex | Tissue | Type | Library | Sequencing/ PCR | Accession No | Comments |
|---|---|---|---|---|---|---|---|---|---|
| *Heliconius charithonia* | Austin, TX | HCH630 | F | Adult | gDNA | Hi-C | | SRR19423652 SRR19423651 | Raw reads |
| *charithonia* | Irvine, CA | HCH2 | F | Pupae | gDNA | PacBio | RS Sequencing | PRJNA505348 | Raw reads |
| *charithonia* | Irvine, CA | HCH2 | F | Pupae | gDNA | PacBio | | PRJNA505348 | genome assembly |
| *charithonia* | Irvine, CA | HCH2 | F | Pupae | gDNA | Illumina | HiSeq 4000 PE150 bp | SRR19659010 SRR19659009 | Raw reads |
| *charithonia* | Irvine, CA | HCH676 | F | Pupae | gDNA | Illumina | HiSeq 4000 PE100 bp | SRR19663609 | Raw reads |
| *charithonia* | Irvine, CA | HCH678 | M | Pupae | gDNA | Illumina | HiSeq 4000 PE100 bp | SRR19663608 | Raw reads |
| *charithonia* | Costa Rica | HCH456 | F | Adult head | RNA | Illumina | PE100 bp | E-MTAB-6810[1] | Raw reads |
| *charithonia* | Costa Rica | HCH457 | F | Adult head | RNA | Illumina | PE100 bp | E-MTAB-6810[1] | Raw reads |
| *charithonia* | Costa Rica | HCH502 | F | Adult head | RNA | Illumina | PE100 bp | E-MTAB-6810[1] | Raw reads |
| *charithonia* | Costa Rica | HCH504 | F | Adult head | RNA | Illumina | PE100 bp | E-MTAB-6810[1] | Raw reads |
| *charithonia* | Costa Rica | HCH506 | F | Adult head | RNA | Illumina | PE100 bp | E-MTAB-6810[1] | Raw reads |
| *charithonia* | Costa Rica | HCH508 | F | Adult head | RNA | Illumina | PE100 bp | E-MTAB-6810[1] | Raw reads |
| *charithonia* | Costa Rica | HCH453 | M | Adult head | RNA | Illumina | PE100 bp | E-MTAB-6810[1] | Raw reads |
| *charithonia* | Costa Rica | HCH454 | M | Adult head | RNA | Illumina | PE100 bp | E-MTAB-6810[1] | Raw reads |
| *charithonia* | Costa Rica | HCH501 | M | Adult head | RNA | Illumina | PE100 bp | E-MTAB-6810[1] | Raw reads |
| *charithonia* | Costa Rica | HCH503 | M | Adult head | RNA | Illumina | PE100 bp | E-MTAB-6810[1] | Raw reads |

| | | | | | | | | | |
|---|---|---|---|---|---|---|---|---|---|
| *charithonia* | Costa Rica | HCH505 | M | Adult head | RNA | Illumina | PE100 bp | E-MTAB-6810[1] | Raw reads |
| *charithonia* | Costa Rica | HCH507 | M | Adult head | RNA | Illumina | PE100 bp | E-MTAB-6810[1] | Raw reads |
| *charithonia* | Costa Rica | HCH708 | F | antennae | RNA | Illumina | PE100 bp | SRR19860620 | Raw reads |
| *charithonia* | Costa Rica | HCH709 | F | antennae | RNA | Illumina | PE100 bp | SRR19860610 | |
| *charithonia* | Costa Rica | HCH770 | F | antennae | RNA | Illumina | PE100 bp | | |
| *charithonia* | Costa Rica | HCH771 | F | antennae | RNA | Illumina | PE100 bp | | |
| *charithonia* | Costa Rica | HCH713 | M | antennae | RNA | Illumina | PE100 bp | SRR19860611 | |
| *charithonia* | Costa Rica | HCH724 | M | antennae | RNA | Illumina | PE100 bp | SRR19860603 | |
| *charithonia* | Costa Rica | HCH765 | M | antennae | RNA | Illumina | PE100 bp | SRR19860613 | |
| *charithonia* | Costa Rica | HCH768 | M | antennae | RNA | Illumina | PE100 bp | SRR19860609 | |
| *charithonia* | Costa Rica | HCH708 | F | mouthparts | RNA | Illumina | PE100 bp | SRR19860608 | |
| *charithonia* | Costa Rica | HCH715 | F | mouthparts | RNA | Illumina | PE100 bp | SRR19860606 | |
| *charithonia* | Costa Rica | HCH783 | F | mouthparts | RNA | Illumina | PE100 bp | | |
| *charithonia* | Costa Rica | HCH785 | F | mouthparts | RNA | Illumina | PE100 bp | | |
| *charithonia* | Costa Rica | HCH713 | M | mouthparts | RNA | Illumina | PE100 bp | SRR19860604 | |
| *charithonia* | Costa Rica | HCH724 | M | mouthparts | RNA | Illumina | PE100 bp | SRR19860615 | |
| *charithonia* | Costa Rica | HCH740 | M | mouthparts | RNA | Illumina | PE100 bp | SRR19860617 | |
| *charithonia* | Costa Rica | HCH741 | M | mouthparts | RNA | Illumina | PE100 bp | SRR19860618 | |
| *charithonia* | Costa Rica | HCH708 | F | legs | RNA | Illumina | PE100 bp | SRR19860619 | |
| *charithonia* | Costa Rica | HCH709 | F | legs | RNA | Illumina | PE100 bp | SRR19860607 | |
| *charithonia* | Costa Rica | HCH770 | F | legs | RNA | Illumina | PE100 bp | | |
| *charithonia* | Costa Rica | HCH771 | F | legs | RNA | Illumina | PE100 bp | | |
| *charithonia* | Costa Rica | HCH713 | M | legs | RNA | Illumina | PE100 bp | SRR19860605 | |
| *charithonia* | Costa Rica | HCH724 | M | legs | RNA | Illumina | PE100 bp | SRR19860616 | |
| *charithonia* | Costa Rica | HCH740 | M | legs | RNA | Illumina | PE100 bp | | |
| *charithonia* | Costa Rica | HCH741 | M | legs | RNA | Illumina | PE100 bp | SRR19860614 | |
| *charithonia* | Costa Rica | | | | | | | | stringtie assembly |
| *charithonia* | Costa Rica | HCH676 | F | Adult | DNA | PCR | Sanger | | |
| *charithonia* | Costa Rica | HCH678 | M | Adult | DNA | PCR | Sanger | | |
| *cydno* | Costa Rica | HCY805 | F | Adult | DNA | PCR | Sanger | | |
| *cydno* | Costa Rica | HCY806 | M | Adult | DNA | PCR | Sanger | | |
| *doris* | Costa Rica | HDO800 | F | Adult | DNA | PCR | Sanger | | |
| *doris* | Costa Rica | HDO803 | M | Adult | DNA | PCR | Sanger | | |
| *erato* | Costa Rica | HER2013 | F | Adult | DNA | PCR | Sanger | | |
| *erato* | Costa Rica | HER2018 | M | Adult | DNA | PCR | Sanger | | |
| *hecale* | Costa Rica | HHE801 | F | Adult | DNA | PCR | Sanger | | |
| *hecale* | Costa Rica | HHE803 | M | Adult | DNA | PCR | Sanger | | |
| *hewitsoni* | Costa Rica | HHW801 | F | Adult | DNA | PCR | Sanger | | |
| *hewitsoni* | Costa Rica | HHW803 | M | Adult | DNA | PCR | Sanger | | |
| *ismenius* | Costa Rica | HIS801 | F | Adult | DNA | PCR | Sanger | | |
| *ismenius* | Costa Rica | HIS803 | M | Adult | DNA | PCR | Sanger | | |
| *melpomene* | Costa Rica | HME1312 | F | Adult | DNA | PCR | Sanger | | |
| *melpomene* | Costa Rica | HME1315 | M | Adult | DNA | PCR | Sanger | | |
| *sara* | Costa Rica | HSA801 | F | Adult | DNA | PCR | Sanger | | |
| *sara* | Costa Rica | HSA804 | M | Adult | DNA | PCR | Sanger | | |
| *sapho* | Costa Rica | HSP801 | F | Adult | DNA | PCR | Sanger | | |
| *sapho* | Costa Rica | HSP803 | M | Adult | DNA | PCR | Sanger | | |

[1]RNA-seq data from Catalan et al. (2018) Evolution of sex-biased gene expression and dosage compensation in the eye and brain of *Heliconius* butterflies. *Mol. Biol. Evol.* 35:2120-2134.
